# AtCGL160 recruits chloroplast coupling factor 1

**DOI:** 10.1101/2021.10.01.462544

**Authors:** Bennet Reiter, Lea Rosenhammer, Giada Marino, Stefan Geimer, Dario Leister, Thilo Rühle

**Affiliations:** Plant Molecular Biology Faculty of Biology I, Ludwig-Maximilians-Universität Munich, D-82152 Planegg-Martinsried, Germany; Zellbiologie/Elektronenmikroskopie NW I/B1, Universität Bayreuth, 95447 Bayreuth, Germany

**Keywords:** chloroplast, photosynthesis, ATP synthase, thylakoid complex, assembly, CF_1_-CF_O_, Arabidopsis

## Abstract

ATP synthases couple the generation of chemical energy to a transmembrane electro-chemical potential. Like ATP synthases in bacteria and mitochondria, chloroplast ATP synthases consist of a membrane-spanning (CF_O_) and a soluble coupling factor (CF_1_). Accessory factors facilitate subunit production and orchestrate the assembly of the functional CF_1_-CF_O_ complex. It was previously shown that the accessory factor CGL160 promotes the formation of plant CF_O_ and performs a similar function in the assembly of its c-ring to that of the distantly related bacterial Atp1/UncI protein. In this study, we show that the N-terminal portion of CGL160 (AtCGL160N), which is specific to the green lineage, is required for late steps in CF_1_-CF_O_ assembly in *Arabidopsis thaliana*. In plants that lacked this stroma-exposed domain, photosynthesis was impaired, and amounts of CF_1_-CF_O_ were reduced to about 65% of the wild-type level. Loss of AtCGL160N did not perturb c-ring formation, but led to a 10-fold increase in the numbers of CF_1_ sub-complexes in the stroma relative to the wild type and the CF_1_ assembly mutant *atcgld11-1*. Co-immunoprecipitation and protein crosslinking assays revealed an association of AtCGL160 with CF_1_ subunits. Yeast two-hybrid assays localized the interaction to a stretch of AtCGL160N that binds to the thylakoid-proximal domain of CF_1_-β that includes the conserved DELSEED motif. We therefore propose that AtCGL160 has acquired an additional function in the recruitment of soluble CF_1_ to a membrane-integral CF_O_ sub-complex, which is critical for the modulation of CF_1_-CF_O_ activity and photosynthesis in chloroplasts.

## Introduction

F-type ATP synthases, which utilize chemiosmotic membrane potentials to generate ATP, are central actors in the energy metabolism of bacteria, mitochondria and chloroplasts. These biological nanomotors share a largely conserved structure, consisting of a soluble F_1_ and a membrane-bound F_O_ moiety. Bacterial and chloroplast ATP synthases (CF_1_-CF_O_) are closely related with respect to size and subunit composition (Groth and Pohl, 2001; Vollmar et al., 2009; Hahn et al., 2018) and, in contrast to the multimeric mitochondrial ATP synthases, exist as monomers in thylakoid membranes (Daum et al., 2010). In the chloroplasts of higher plants, CF_1_-CF_O_ complexes reside exclusively in stroma lamellae and grana-end membranes, because the ~16-nm stromal extension of CF_1_ prevents its incorporation into the tightly packed grana stacks (Daum et al., 2010).

During photophosphorylation, CF_1_-CF_O_ complexes couple the light-driven generation of the trans-thylakoid proton-motive force (*pmf*) to ADP phosphorylation. The membrane-embedded proteolipidic c_14_-ring, together with the non-covalently bound central stalk γε, form the motor unit, and drive rotary catalysis by CF_1_. The peripheral stator consists of the subunits a, b and b’, and is connected to the (αβ)_3_ unit by the δ subunit, which acts as a flexible hinge between CF_1_ and CF_O_ (Murphy et al., 2019). Protons are translocated from the luminal to the stromal side via two aqueous channels in the a subunit. During translocation, each proton enters the access channel and binds to a conserved glutamate residue in subunit c. The c_14_ motor executes an almost complete rotation before releasing the proton into the stroma through the exit channel (Hahn et al., 2018). The counterclockwise rotation of the central stalk in the vicinity of the hexamer triggers alternating nucleotide-binding affinities in the β subunits that ultimately drive ATP generation (reviewed in von Ballmoos et al., 2009; Junge and Nelson, 2015).

As a result of extensive organellar gene transfer during plant evolution, three CF_1_-CF_O_ subunits (b’, γ, δ) are encoded in the nuclear genome, while the remaining CF_1_-CF_O_ genes are organized into two plastid operons. Consequently, two different gene-expression systems must be tightly coordinated with the chloroplast protein import machinery for efficient CF_1_-CF_O_ biogenesis. Several CF_1_-CF_O_ auxiliary factors involved in plastid gene expression have been identified, including proteins involved in mRNA processing (AEF1), mRNA stabilization (PPR10, BFA2) and translation initiation (ATP4, TDA1) (Pfalz et al., 2009; Eberhard et al., 2011; Zoschke et al., 2012; Yap et al., 2015; Zhang et al., 2019). Moreover, CF_1_-CF_O_ assembly factors ensure correct complex stoichiometry, and prevent the accumulation of dead-end products or harmful intermediates that could lead to wasteful ATP hydrolysis or *pmf* dissipation.

As in the case of the bacterial assembly model, plastid CF_1_-CF_O_ complexes are constructed from different intermediates or modules (reviewed in Rühle and Leister, 2015). CF_1_ assembly was first examined using in-vitro reconstitution assays, and was shown to be initiated by α/β dimerization in a chaperone-assisted process (Chen and Jagendorf, 1994). CF_1_ formation depends on CGLD11/BFA3, which is specific to green plants, interacts with the hydrophobic catalytic site of the β-subunit and may prevent aggregation or formation of unfavorable homodimers (Grahl et al., 2016; Zhang et al., 2016). Moreover, PAB (Mao et al., 2015) and BFA1 (Zhang et al., 2018) have been proposed to be required for efficient incorporation of the γ subunit into CF_1_.

Less is known about CF_O_ assembly, and only one accessory factor – CONSERVED ONLY IN THE GREEN LINEAGE 160 (CGL160) – has been identified so far (Rühle et al., 2014). Absence of CGL160 in the *Arabidopsis thaliana* mutant *atcgl160-1* is associated with a significant reduction (70-90%) in wild-type CF_1_-CF_O_ levels, and CF_O_-c subunits accumulate as monomers. Moreover, split-ubiquitin assays have provided evidence that AtCGL160 interacts with CF_O_-c and CF_O_-b. It was therefore concluded that AtCGL160 is required for efficient formation of the c-ring in chloroplasts and shares this function with its distantly related bacterial counterpart Atp1/UncI (Suzuki et al., 2007; Ozaki et al., 2008). Furthermore, AtCGL160 was suggested to participate in CF_1_ assembly into the holo-complex, based on CF_1_ subcomplex co-migration and crosslinking experiments using a putatively specific anti-AtCGL160 antibody (Fristedt et al., 2015).

In this study, the function of the N-terminal domain that is conserved in all CGL160 proteins from the green lineage was investigated in Arabidopsis. The results demonstrate that this domain (AtCGL160N) mediates the critical connection of CF_1_ to CF_O_ assembly modules by interacting with subunit β. Thus, CGL160 emerges as a key auxiliary factor that not only promotes CF_O_ formation, but is also involved in late CF_1_-CF_O_ assembly steps.

## Results

### The N-terminal moiety of AtCGL160 is required for efficient photosynthesis and CF_1_-CF_O_ functionality

CGL160 was identified based on its coregulation with photosynthetic genes in ATTED-II (Obayashi et al., 2009) and its affiliation to the GreenCut suite of proteins (Merchant et al., 2007; Karpowicz et al., 2011). The C-terminal transmembrane segment of CGL160 (~15 kDa) is distantly related to bacterial Atp1/UncI (Rühle et al., 2014; Fristedt et al., 2015), whereas the larger N-terminal portion of the protein sequence is only conserved in algae, bryophytes, and higher plants (Supplemental Fig. 1). This latter domain of ~200 amino acids (aa) in *Arabidopsis thaliana* (AtCGL160N) includes a predicted N-terminal chloroplast transit peptide (cTP) of 46 aa (Emanuelsson et al., 1999), and mass spectrometry has identified several phosphorylated peptides which are derived from positions 106-134 (Reiland et al., 2009; Reiland et al., 2011; Roitinger et al., 2015). Indeed, two conserved putative phosphorylation sites were found in a multiple sequence alignment of CGL160 homologs from species across the green lineage, which correspond to positions S111 and S126 in AtCGL160 (Fig. 1, Supplemental Fig. 1).

**Figure 1.**
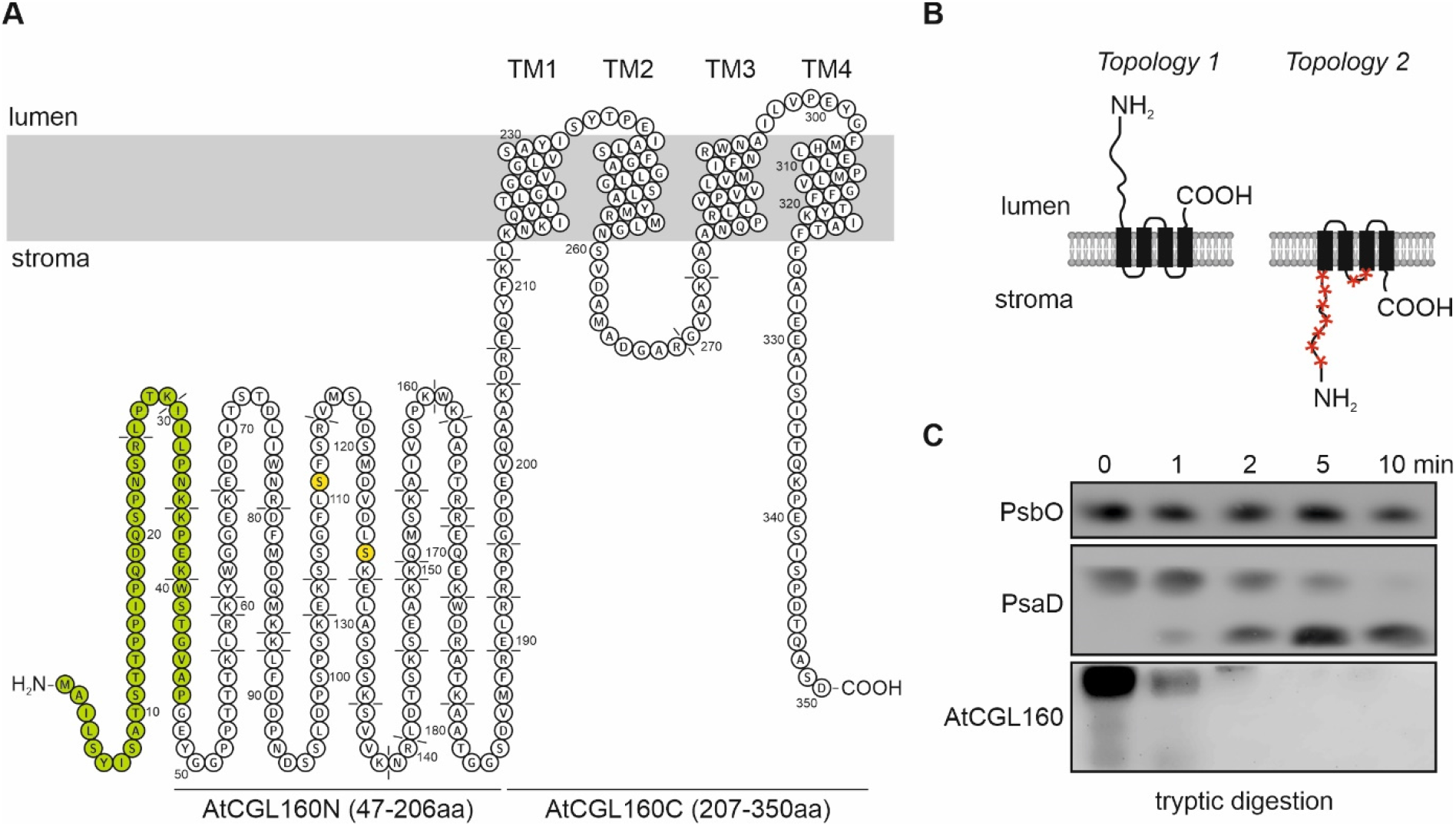
Topology of AtCGL160 and trypsin cleavage-site prediction. **A,** Transmembrane (TM) domain predictions were obtained from the AtCGL160 UniProt protein accession O82279. Putative trypsin cleavage sites are highlighted in dashed lines and amino-acid positions are indicated. The topology was drawn for the full-length sequence of AtCGL160 including the predicted transit peptide (green) with Protter (Omasits et al., 2014). Two conserved serine residues (S111 and S126) are marked in yellow. **B,** Representation of two putative AtCGL160 topologies. The four transmembrane domains are indicated as black boxes. Accessible trypsin digestion sites are highlighted by red stars. **C,** Immunoblot of thylakoid membranes of the WT (Col-0) fractionated by SDS-PAGE, untreated (0 min) or treated with trypsin for 1, 2, 5 and 10 min. Blots were probed with antibodies against the lumen-oriented PSII subunit PsbO, the stroma-exposed PSI subunit PsaD and AtCGL160.

Earlier studies have provided experimental evidence for the localization of AtCGL160 to the thylakoid membrane (Rühle et al., 2014; Tomizioli et al., 2014; Fristedt et al., 2015). To gain further insights into the topology of AtCGL160, a protease protection assay was carried out (Fig. 1B, C). In the case of topology 1, all trypsin cleavage sites in AtCGL160 reside in the lumen of the thylakoid and remain fully protected from proteolytic degradation (Fig. 1B). Conversely, the stromal orientation of AtCGL160N predicted for topology 2 would expose trypsin cleavage sites and lead to degradation products of less than 2 kD (Fig. 1A). To test the accessibility of native AtCGL160N, wild-type thylakoids were isolated and treated with trypsin for 10 min (Fig. 6B). As expected, the luminal PSII subunit PsbO was not affected by the enzyme, whereas the stromally exposed PSI subunit PsaD was susceptible to the protease. AtCGL160N was also efficiently digested, leaving no detectable proteolytic cleavage products, which is consistent with protrusion of the entire N-terminal domain into the stroma, as shown in topology 2 (Fig. 6A,B).

To dissect the function of the N-terminal portion of AtCGL160, three different constructs under control of the 35S promoter were cloned, transformed into the *atcgl160-1* background and screened for complementation (Fig. 2, Supplemental Fig. 2A). Plants that overexpressed the full-length coding sequence (CDS) of *AtCGL160* served as controls (*P_35S_:AtCGL160*), while the other two genotypes expressed either the CDS of the N-terminal (*P_35S_:AtCGL160N*) or the C-terminal segment (*P_35S_:AtCGL160C*) of the protein (Supplemental Fig. 2B, C). In the case of *P_35S_:AtCGL160C* plants, targeting of the truncated version to chloroplasts was achieved by fusing the CDS of the AtCGL160-derived cTP (1-46 aa) to that of AtCGL160C (Fig. 2A). As was previously demonstrated in complementation analyses with *P_35S_:AtCGL160-eGFP* lines (Rühle et al., 2014), overexpression of the full-length *AtCGL160* rescued the *atcgl160-1* phenotype (Supplemental Fig. 2A), as indicated by wild-type-like growth and restored leaf morphology under short-day conditions (Fig. 2B, C). *P_35S_:AtCGL160N* failed to complement the mutant phenotype (Fig. 2B, C, Supplemental Fig. 2A) and AtCGL160N could not be detected in either stromal or thylakoid extracts (Supplemental Fig. 3). Since *AtCGL160N* transcripts were present in WT-like amounts in *P_35S_:AtCGL160N* plants (Supplemental Fig. 2C), the lack of AtCGL160N is probably due to proteolytic degradation owing to its inability to associate correctly with thylakoids. Nevertheless, *P_35S_:AtCGL160N* plants were retained and served as an additional AtCGL160 knockout control. *P_35S_:AtCGL160C* plants with similar overexpression rates to *P_35S_:AtCGL160* plants (Supplemental Fig. 2B, C) were characterized by a significant increase in leaf area compared to the mutant background *atcgl160-1*, but were growth-retarded with respect to the wild-type control. Interestingly, like *atcgl160-1*, *P_35S_:AtCGL160C* plants developed a variegated phenotype in old leaves, which was not found either in the wild type or in the CF_1_ assembly mutant *atcgld11-1* (Grahl et al., 2016) under short-day conditions (Fig. 2B).

**Figure 2.**
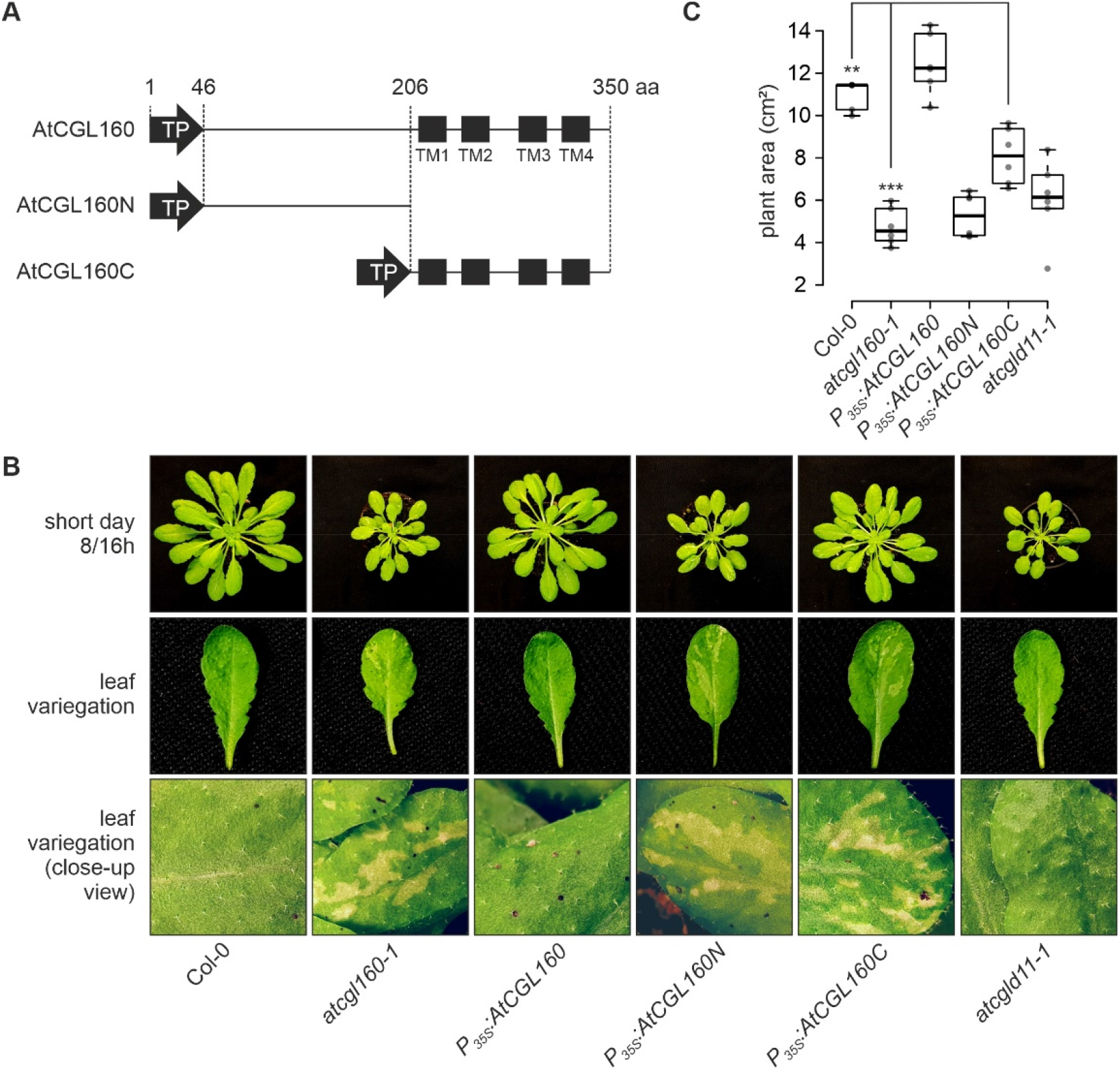
Growth phenotype and leaf variegation of *P_35S_:AtCGL160, P_35S_:AtCGL160N* and *P_35S_:AtCGL160C* plants under short-day conditions. **A**, Schematic representations of reintroduced AtCGL160 coding sequences. Plants lacking AtCGL160 were transformed with overexpressor constructs harboring the coding sequences for the full-length AtCGL160 (*P_35S_:AtCGL160*) and its N- (*P_35S_:AtCGL160N*) and C-terminal (*P_35S_:AtCGL160C*) segments. Transcription was under the control of the 35S CaMV promoter and targeting to the chloroplast was mediated by the transit peptide of AtCGL160 (TP). Amino-acid positions are indicated and predicted transmembrane domains (TM1-TM4) are schematically shown as black boxes. **B**, Leaf morphology of Col-0, *atcgl160-1*, *P_35S_:AtCGL160, P_35S_:AtCGL160N*, *P_35S_:AtCGL160C* and *atcgld11-1* plants. **C**, Leaf areas of 6 individual plants per genotype were determined 4 weeks after germination. The horizontal lines represent the median, and boxes indicate the 25th and 75th percentiles. Whiskers extend the interquartile range by a factor of 1.5×, and outliers are represented by dots. The effect of the deletion of AtCGL160N in *P_35S_:AtCGL160C* plants on growth under short-day conditions was tested by paired sample t-test (two-sided). Statistically significant differences are marked with asterisks (\**P*<0.05, \*\**P*<0.01, and \*\*\**P*<0.001).

To analyze the leaf phenotype in more detail, we carried out electron microscopic analyses of Col-0, *atcgl160-1* and *P_35S_:AtCGL160C* plants (Fig. 3). In these genotypes, the chloroplast ultrastructure in preparations from green leaf sections was unchanged with regard to thylakoid content, curvature and grana organization (Fig. 3 A-F). These observations in *atcgl160-1*, together with previous ultrastructural analyses of the CF_1_ assembly mutant line *atcgld11-1* (Grahl et al., 2016) and spinach chloroplasts (Daum et al., 2010), support the idea that CF_1_-CF_O_ complexes are not physically involved in thylakoid curvature formation. Examination of white leaf sections in *atcgl160-1* and *P_35S_:AtCGL160C* revealed the absence of thylakoids in plastids, accompanied by the appearance of plastoglobuli in densely packed stromal clusters (Fig. 3 G-J). Furthermore, large vesicles were observed, which also point to increased catabolic activity and degradation processes in *atcgl160-1* and *P_35S_:AtCGL160C* plastids. Another finding was the inclusion of mitochondria in degraded plastids, which was also observed, to a lesser extent, in white leaf sectors of *P_35S_:AtCGL160C*.

**Figure 3.**
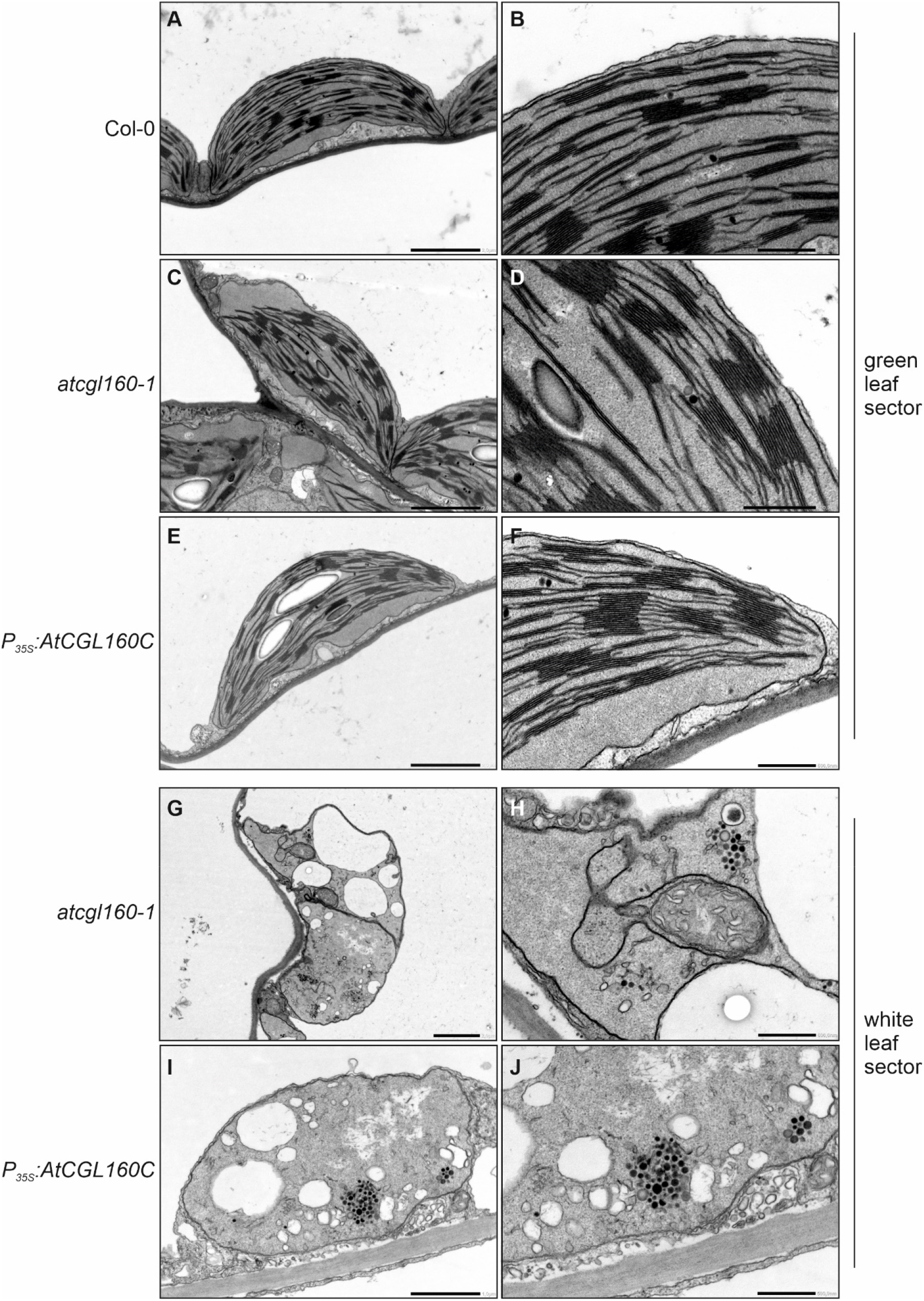
Plastid ultrastructure in white leaf sectors is altered in the absence of AtCGL160N under short-day growth conditions. Electron micrographs of samples from green leaf sections obtained from Col-0 **(A, B)**, *atclg160-1* **(C, D)** and *P_35S_:AtCGL160C* **(E, F)** plants. The ultrastructure of chloroplasts was further examined in samples of white leaf sections obtained from *atcgl160-1* **(G, H)** and *P_35S_:AtCGL160C* **(I, J)** plants. The photos on the right show enlargements of the images on the left. The scale bar corresponds to 2 µm in **A**, **C**, **E**, and **G**, 1 µm in **I** and 0.5 µm in **B**, **D**, **F**, **H** and **J**.

To test whether disruption of AtCGL160N impairs photosynthesis and CF_1_-CF_O_ activity, measurements of chlorophyll *a* fluorescence and electrochromic shift (ECS) were carried out on Col-0, *atcgl160-1*, *P_35S_:AtCGL160, P_35S_:AtCGL160N*, *P_35S_:AtCGL160C* and *atcgld11-1* plants (Fig. 4A-C). As expected, the CF_1_-CF_O_ assembly mutants *atcgl160-1* and *atcgld11-1* showed higher heat dissipation (indicated as non-photochemical quenching, NPQ) and increased proton-motive force (*pmf*), but lower proton conductivity (gH^+^) through the thylakoid membrane compared to the wild-type control. *P_35S_:AtCGL160* and *P_35S_:AtCGL160N* plants displayed similar levels of NPQ, *pmf* and gH^+^ to the wild type and the CF_1_-CF_O_ assembly mutant *atcgld11-1*, respectively. Notably, photosynthetic parameters were only partially restored in *P_35S_:AtCGL160C* lines.

**Figure 4.**
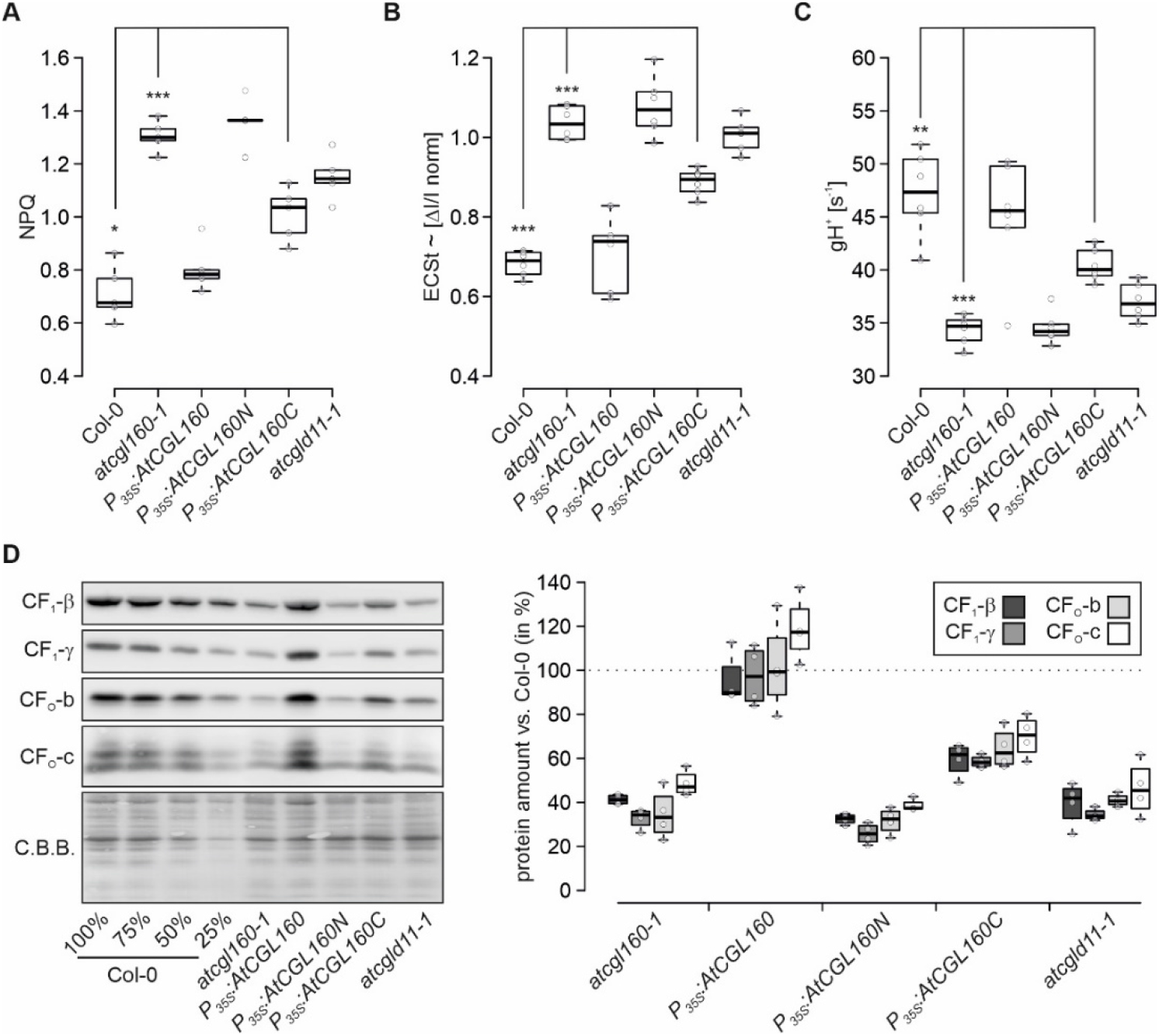
Lack of AtCGL160N perturbs photosynthesis and CF_1_-CF_O_ integrity>. **A**, Heat dissipation (non-photochemical quenching, NPQ) in Col-0, *atcgl160-1*, *P_35S_:AtCGL160, P_35S_:AtCGL160N*, *P_35S_:AtCGL160C* and *atcgld11-1* plants grown under short-day conditions. NPQ values from five plants per genotype were determined 105 s after light induction (145 µmol photons m^-2^ s^-1^) using an Imaging-PAM system (Walz). **B**, Dark-interval relaxation kinetics (DIRK) derived from ECS signals were recorded after 10 min of illumination from six individual plants grown under short-day conditions. Total amplitude of the P515 differential absorption signal was normalized to a single turnover flash 4 min after the ECS measurement. **C**, Proton conductivity of the thylakoid membrane was determined from ECS signal relaxation rates, which were fitted to a first-order decay function. Calculated rate constants were expressed as gH^+^ [s^-1^]. **D**, Steady-state levels of immunodetected CF_1_-CF_O_ marker subunits. After fractionation of thylakoid proteins on SDS-PAGE and transfer to PVDF membranes, blots were probed with antibodies against CF_1_-β, CF_1_-γ, CF_O_-b, and CF_O_-c. Coomassie Brilliant Blue (C.B.B.) staining is shown as loading control. For quantification, signals from four technical replicates of each marker subunit were normalized to signals detected in Col-0 samples. Horizontal lines represent the median, and boxes indicate the 25th and 75th percentiles. Whiskers extend the interquartile range by 1.5×. The effect of the deletion of AtCGL160N on photosynthetic parameters of *P_35S_:AtCGL160C* plants shown in panels **A-C** was tested in paired-sample t-tests (two-sided). Statistically significant differences are marked with asterisks (\**P*<0.05, \*\**P*<0.01, and \*\*\**P*<0.001).

To assess the integrity of the CF_1_-CF_O_ complex in thylakoids, marker subunits were immunodetected in *atcgl160-1*, *P_35S_:AtCGL160, P_35S_:AtCGL160N*, *P_35S_:AtCGL160C* and *atcgld11-1* plants, and quantified relative to Col-0 samples (Fig. 4D). Levels of CF_1_-β, CF_1_-γ, CF_O_-b and CF_O_-c were restored to normal in *P_35S_:AtCGL160*, but reduced to about 60-65% of wild-type amounts in *P_35S_:AtCGL160C* plants. Transformation with the *P_35S_:AtCGL160N* construct had no effect on CF_1_-CF_O_ subunit levels in the *atcgl160-1* mutant.

Overall, overexpression of the Atp1/Unc1-like AtCGL160 domain alone (AtCGL160C) in the *atcgl160-1* background only partially restored CF_1_-CF_O_ amounts (Fig. 4D) and activity (Fig. 4C). Consequently, ∆pH-dependent quenching mechanisms (Fig. 4A) were more highly activated, resulting in downregulation of photosynthesis (Fig. 4B) and growth impairment of *P_35S_:AtCGL160C* plants. We deduced from these results that AtCGL160N might also be involved in CF_1_-CF_O_ assembly at steps other than CF_O_-c ring formation.

### Stromal accumulation of CF_1_ in the absence of AtCGL160N

To investigate the effects of deletion of AtCGL160N on CF_1_-CF_O_ assembly, we performed BN/SDS-PAGE (2D-PAGE) analysis on thylakoids isolated from *P_35S_:AtCGL160* and *P_35S_:AtCGL160C* plants grown under short-day conditions. Consistent with the accumulation of CF_1_-CF_O_ marker subunits in Fig. 4D, CF_1_-β, CF_O_-b and CF_O_-c levels were reduced in *P_35S_:AtCGL160C* compared to plants that overexpressed the full-length CDS of AtCGL160 (Fig. 5A). No accumulation of pre-complexes was observed, as amounts of free proteins, and components of the c-ring, CF_1_ and the holo-complex were reduced uniformly. To assess the assembly status of the c-ring in more detail, we carried out 2D-PAGE with increased amounts of *atcgl160-1* and *P_35S_:AtCGL160C* thylakoids (Fig. 5B). C-ring levels were considerably higher in *P_35S_:AtCGL160C* than in the *atcgl160-1* mutant background. We also examined CF_1_ accumulation in the stroma of Col-0, *P_35S_:AtCGL160, P_35S_:AtCGL160N*, *P_35S_:AtCGL160C* and *atcgld11-1* plants (Fig. 5C), since CF_1_-CF_O_ assembly takes place in a modular fashion and involves distinct thylakoid-integral and soluble intermediates. Strikingly, CF_1_-β and CF_1_-γ were enriched about 10-fold in the stroma of *atcgl160-1*, *P_35S_:AtCGL160N* and *P_35S_:AtCGL160C*, but were detected in close to wild-type levels in *P_35S_:AtCGL160* and *atcgld11-1* plants. In-depth 2D-PAGE analysis of CF_1_ intermediates in *atcgl160-1*, and comparison with results from the co-migration database for photosynthetic organisms (PCom-DB, Takabayashi et al., 2017), revealed that in *atcgl160-1* stromal CF_1_-β and CF_1_-γ were predominantly present in an α_3_β_3_γε complex that lacked subunit CF_1_-δ (Supplemental Fig. S4).

**Figure 5.**
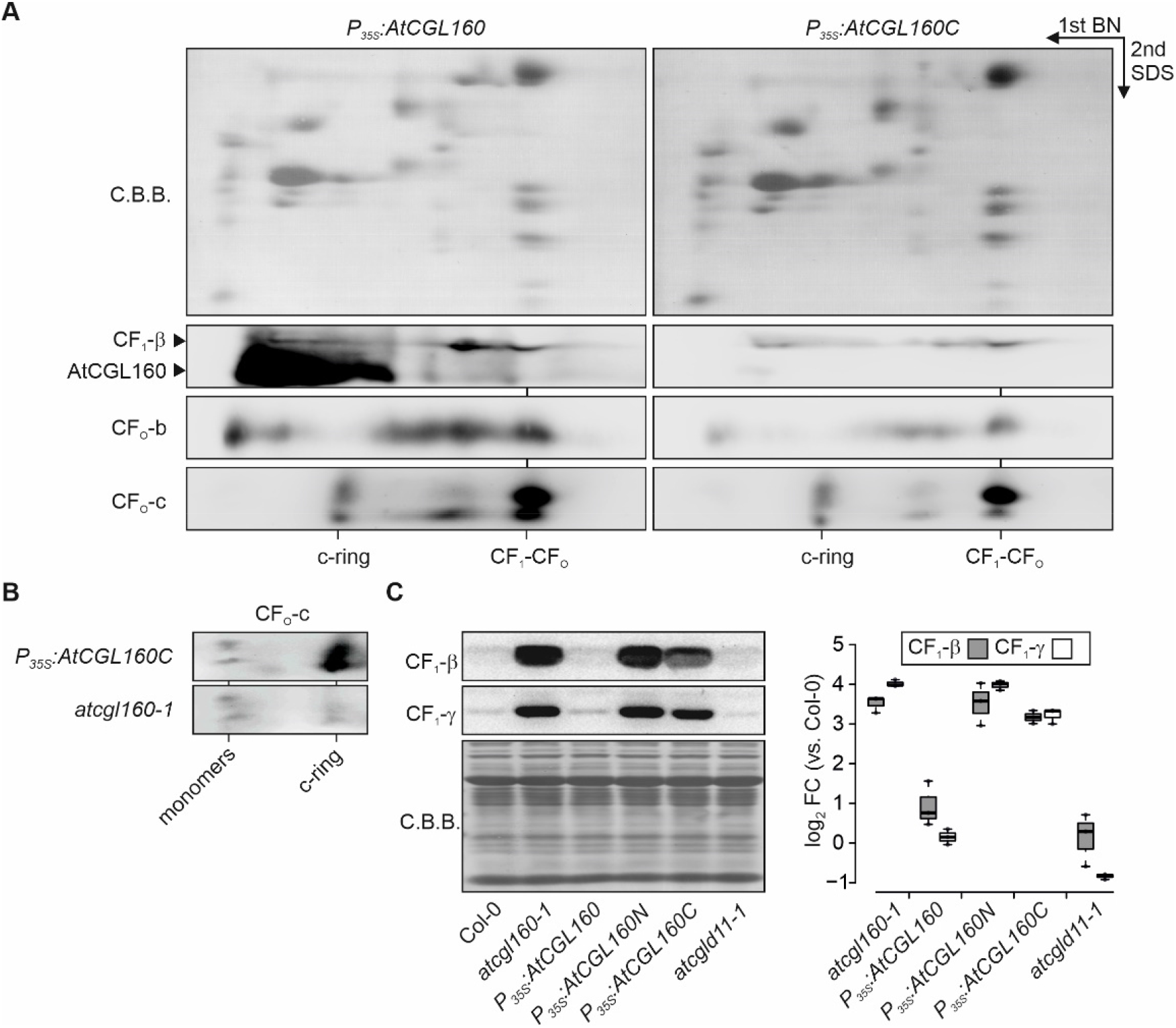
CF_1_-CF_O_ assembly is perturbed in the absence of AtCGL160N. **A**, Thylakoid complexes from *P_35S_:AtCGL160* and *P_35S_:AtCGL160C* plants were solubilized with *n*-dodecyl *β*-D-maltoside (1% [w/v]) and further separated by Blue-Native (BN, 1^st^ dimension) and SDS-PAGE (SDS, 2^nd^ dimension). After protein transfer, PVDF membranes were probed with antibodies against CF_1_-β, CF_O_-b and CF_O_-c, and CF_1_-β blots were subsequently exposed to anti-AtCGL160 antibodies. Positions of the ATP synthase holo-complex (CF_1_-CF_O_) and the c-ring are indicated. Coomassie Brilliant Blue G-250 (C.B.B.) staining of PVDF membranes is shown as loading control. **B**, C-ring assembly in *atcgl160-1* and *P_35S_:AtCGL160C* plants. Increased amounts of thylakoid complexes (corresponding to 120 µg total chlorophyll) were solubilized and fractionated by BN/SDS-PAGE. Blots were probed with an antibody against CF_O_-c. Positions of free c-monomers and the assembled c-ring are indicated. **C**, CF_1_-β and CF_1_-γ enrichment in stromal extract, which was isolated from Col-0, *atcgl160-1*, *P_35S_:AtCGL160, P_35S_:AtCGL160N*, *P_35S_:AtCGL160C* and *atcgld11-1* rosette leaves. Signals of three CF_1_-β and CF_1_-γ immunodetection assays were quantified and are shown on a logarithmic scale. Horizontal lines represent the median, boxes indicate the 25th and 75th percentiles and whiskers extend the interquartile range by a factor of 1.5×.

We concluded that re-introduction of the transmembrane Atp1/Unc1-like domain of AtCGL160 restores c-ring formation, but leads to an overall reduction in CF_1_-CF_O_ levels due to a defect in the attachment of CF_1_ to a membrane-integral CF_O_ intermediate.

### AtCGL160 interacts physically with CF_1_-containing complexes

To pinpoint the role of AtCGL160 in the recruitment of CF_1_ to a membrane-integral CF_O_ intermediate, protein interactions were assessed in co-immunoprecipitation (co-IP) assays (Fig. 6B). Quantitative data for precipitated proteins were obtained by tryptic digestion and subsequent peptide-fragment analysis using liquid chromatography coupled to mass spectrometry. Since the commercially available AtCGL160 antibody (Agrisera AS12 1853) displayed non-specific binding to either CF_1_-α or CF_1_-β (Supplemental Fig. 3), an AtCGL160 antibody with no significant cross-reactions to other thylakoid proteins was generated (Supplemental Fig. 3). In the first step of antibody production, the N-terminal part of AtCGL160 (AtCGL160_29-206aa_) was fused to the maltose-binding protein and injected into rabbits. In the second step, antibodies specific for AtCGL160_29-206aa_ were affinity-purified from rabbit antisera using an immobilized fusion protein consisting of AtCGL160_29-206aa_ and glutathione S-transferase. As expected, when the resulting antibody fraction was tested in immunodetection assays, it showed only one distinct signal in the WT sample, which was enriched in extracts of *P_35S_:AtCGL160*, but was absent in both the *atcgl160-1* mutant and in *P_35S_:AtCGL160C* samples (Supplemental Fig. 3).

**Figure 6.**
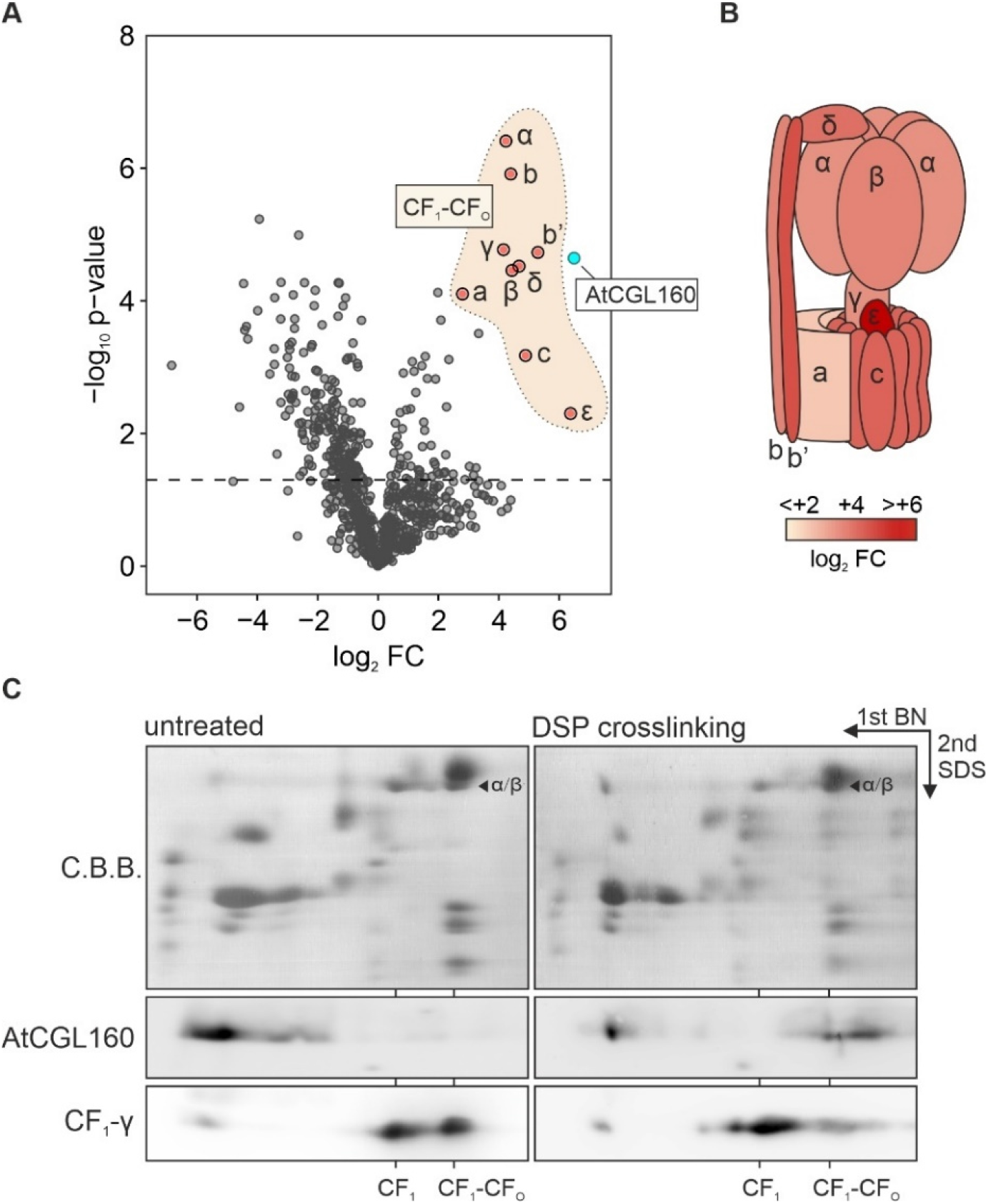
AtCGL160 association with CF_1_ subunits. **A,** Co-immunoprecipitation analyses were carried out with solubilized thylakoids isolated from *P_35S_:AtCGL160*, while *P_35S_:AtCGL160C* plants served as the negative control. Co-immunoprecipitated proteins were further subjected to tryptic digestion, and peptides were analyzed by liquid chromatography coupled to mass spectrometry. Data for differentially enriched proteins are presented in a volcano plot. The relative abundance (log_2_ fold change [log_2_ FC]) of proteins co-immunoprecipitated from *P_35S_:AtCGL160* samples is plotted against their statistically significant enrichment as Benjamini-Hochberg corrected p-values (-log_10_ p-value). The dashed line indicates a negative log_10_ p-value of 1.5, and was defined as the threshold for robust reliability of differences in co-immunoprecipitation data. Blue and red dots highlight quantification results for AtCGL160 and CF_1_-CF_O_ subunits, respectively. **B**, Schematic representation of differentially enriched subunits in a CF_1_-CF_O_ cartoon. Relative amounts of co-immunoprecipitated CF_1_-CF_O_ subunits are shown in colors on a log_2_ FC scale from white (log_2_ FC < 2) to red (log_2_ FC > 6). Co-immunoprecipitation assays were carried out on three independent biological replicates. **C,** Co-migration of AtCGL160 with CF_1_-CF_O_ in crosslinking experiments. Two-dimensional BN/SDS-PAGE analysis was used to compare untreated thylakoid extracts of the WT (Col-0) with extracts that had been crosslinked with dithiobis(succinimidyl propionate) (DSP). Blots of the second dimension were probed with antibodies against AtCGL160 and CF_O_-γ. The positions of CF_1_-CF_O_, the CF_1_ intermediate, and the free protein fraction are indicated based on the mobility of α/β on the C.B.B. stained gel.

Next, NP40-solubilized thylakoid proteins from *P_35S_:AtCGL160* plants grown under short-day conditions were chosen as co-IP input and pulled-down protein amounts were compared to those recovered in co-IP experiments carried out on thylakoid protein extracts of *P_35S_:AtCGL160C*. Plants devoid of AtCGL160 were not considered for use as negative controls, since the reduction in CF_1_-CF_O_ levels observed in *atcgl160-1* (and *P_35S_:AtCGL160N*) (Fig. 4D) might lead to misinterpretation of differential co-IP experiments. As expected, AtCGL160 was pulled down efficiently from *P_35S_:AtCGL160C* extracts (log_2_ FC ~6.5). Moreover, all CF_1_-CF_O_ subunits were identified in co-IPs (Fig. 6A,B) with high differential enrichment levels for the subunits α, β, γ, δ, ε, b, b’ and c (log_2_ FC > 4.4). Subunit CF_O_-a was co-immunoprecipitated at lower levels (log_2_ FC ~2.8). The pull-down of CF_1_ subunits was confirmed by immunodetection assays of the two marker subunits CF_1_-β and CF_1_-γ, which were only detectable in co-IP output fractions obtained from *P_35S_:AtCGL160* samples (Supplemental Fig. 5). Other known CF_1_-CF_O_ assembly factors were not co-immunoprecipitated (Supplemental Table 3), indicating that AtCGL160 is associated with a late CF_1_-CF_O_ assembly stage or the fully assembled complex from which other auxiliary factors had already dissociated.

To confirm the association of AtCGL160 with CF_1_-containing complexes, crosslinking experiments were also carried out (Fig. 6C). To this end, thylakoid membranes of wild-type plants were treated with the crosslinker dithiobis(succinimidyl propionate) (DSP), and subsequently subjected to 2D-PAGE and immunodetection of AtCGL160 and CF_1_-CF_O_ marker subunits. In analyses with untreated thylakoid samples, AtCGL160 migrated predominantly in the monomer fraction. After crosslinking, AtCGL160 could be detected at a molecular mass range which corresponded to that of the CF_1_-CF_O_ holo-complex.

In summary, co-IP of all CF_1_-CF_O_ subunits with an AtCGL160-specific antibody, together with the observation that AtCGL160 co-migrated with the CF_1_-CF_O_ holo-complex after DSP cross-linking, corroborates the involvement of AtCGL160 in the functional integration of CF_1_ into the holo-complex at a late step in CF_1_-CF_O_ assembly.

### AtCGL160N interacts with CF_1_-β in yeast two-hybrid assays

Interactions between the stroma-oriented AtCGL160N domain and individual CF_1_ subunits were further examined by yeast two-hybrid experiments (Fig. 7). A construct coding for a fusion of AtCGL160N_29-206aa_ to the GAL4-binding domain (BD) was co-transformed into yeast cells together with constructs coding for GAL4 activation domain (AD) fusions to all CF_1_ subunits (α, β, γ, δ, ε). Moreover, BD-AtCGL160N interaction was tested with AD fusions to the soluble parts of the stator subunits b and b’, AtCGL160N, and CF_1_ assembly factor AtCGLD11. As a result, only yeast cells carrying constructs for AD-CF_1_-β and BD-AtCGL160N could grow on selective medium (Fig. 7A). To narrow down the CF_1_-β interaction site, additional AD fusion constructs were cloned that encoded three different CF_1_-β subdomains (Fig. 7B) defined in earlier studies (Groth and Pohl, 2001; Zhang et al., 2016). Domain I comprises a thylakoid-distal β-barrel structure and interacts with CF_1_-δ. Domain II harbors the catalytic site involved in ATP generation or hydrolysis. The thylakoid-proximal domain III contains the conserved “DELSEED” motif, which is required for CF_1_-γ/ε-dependent regulation of ATP hydrolysis and synthase activity (Kanazawa et al., 2017; Hahn et al., 2018). When tested on restrictive medium, only cells harboring AD-CF_1_-β_III_ together with BD-AtCGL160N could grow. In a reciprocal approach, coding sequences of AtCGL160N were deleted successively from the BD-AtCGL160N construct (∆29-74, ∆75-105, ∆106-134, ∆135-160 and ∆161-206 aa) and tested for AD-CF_1_-β interaction in yeast cells (Fig. 7C). Only the ∆29-74 and ∆75-105 deletions resulted in an absence of growth, while yeast strains with deletion constructs of ∆106-134, ∆135-160 and ∆161-206 aa were able to proliferate on selective medium (Fig. 6B). Thus, the interaction between AtCGL160 and CF_1_ involves AtCGL160_29-105_ and the thylakoid-proximal domain of CF_1_-β_III_, while the phosphorylation hotspot identified in the protein segment 106-134 aa (Fig. 1A) is dispensable for the interaction.

**Figure 7.**
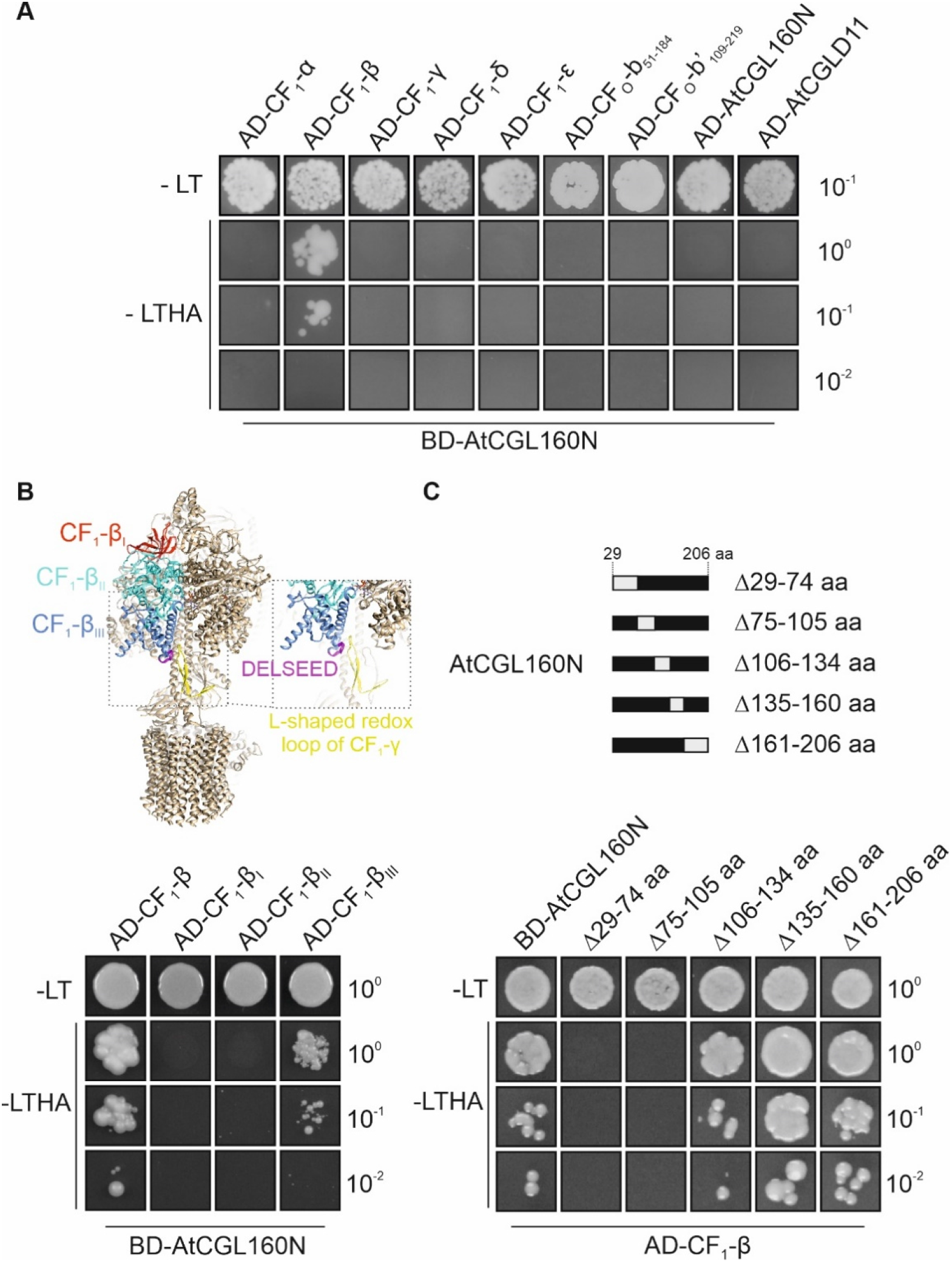
AtCGL160N interaction studies in yeast two-hybrid assays. **A**, Interactions of AtCGL160 with CF_1_-CF_O_ structural components exposed on the stromal side of thylakoids were tested by transformation of a construct that fuses AtCGL160N to the GAL4 DNA-binding domain (BD-AtCGL160). Cells were then co-transformed with constructs coding for GAL4 activation domain (AD) fused to CF_1_-α, β, γ, δ, ε, CF_O_-b_51-184_ or CF_O_-b‘_109-219_, as well as AtCGL160N or AtCGLD11. **B**, Interaction of AtCGL160N with structural domains of CF_1_-β. Yeast cells carrying a construct coding for BD-AtCGL160 were transformed with constructs coding for AD-CF_1_-β_I_, AD-CF_1_-β_II_, and AD-CF_1_-β_III_. Structural domains of the CF_1_-β are colored in red (domain I), turquoise (domain II), and blue (domain III). The conserved DELSEED motif is shown in purple, and the L-shaped redox loop of CF_1_-γ in yellow. The atomic model of CF_1_-CF_O_ was obtained from the PDB database (ID: 6fkh, Hahn et al. (2018)) and formatted with ChimeraX (Pettersen et al., 2021). **C,** Mapping of the AtCGL160N interaction site. Consecutive regions (grey boxes) coding for segments of the soluble AtCGL160 domain were omitted from the BD-AtCGL160N vector and co-transformed with AD-CF_1_-β into competent yeast cells. Transformations were verified by plating on permissive medium lacking Leu and Trp (-LT). Interactions were then tested on selective medium (-Leu/-Trp/-His/-Ade, [-LTHA]) by plating equal numbers of yeast cells in serial dilutions (10^0^, 10^−1^, and 10^−2^).

## Discussion

### AtCGL160N recruits a stromal α3β3γε complex for late CF_1_-CF_O_ assembly steps

Despite structural similarities and comparable subunit compositions, the number of known assembly factors for ATP synthases is markedly higher in chloroplast than in bacterial systems (reviewed in Zhang et al., 2020). Moreover, in plants the Atp1/UncI-related CGL160 assembly factor has acquired an N-terminal domain that is specific for the green lineage. Thus, the expanded molecular inventory for CF_1_-CF_O_ assembly in chloroplasts might reflect the need for tight post-translational control of CF_1_-CF_O_ formation, since the complex plays a central role in *pmf* utilization and regulation of photosynthesis (reviewed in Avenson et al., 2005). In this context, an important finding of previous studies was that disruption of full-length AtCGL160 (Rühle et al., 2014; Fristedt et al., 2015) was more detrimental to levels of functional ATP synthase than the loss of Atp1/UncI in bacteria (Gay, 1984; Liu et al., 2013). Furthermore, we show here that expression of *P35S:AtCGL160C* in plants that lack AtCGL160N only partially restores CF_1_-CF_O_ levels and activity (Fig. 4). These observations prompted us to investigate the molecular function of the green-lineage-specific AtCGL160N in the CF_1_-CF_O_ assembly process in more detail.

Several lines of evidence suggest that the N-terminal domain of AtCGL160 recruits a stromal CF_1_ intermediate, while the C-terminal segment participates in c_14_-ring assembly: (i) AtCGL160N protrudes into the stroma, as deduced from protease protection assays (Fig. 1); (ii) formation of the c_14_ ring is restored in the presence of AtCGL160C alone, but CF_1_ accumulates strongly in the stroma in the absence of AtCGL160N (Fig. 5), (iii) CF_1_ subunits are differentially enriched in co-IP analyses performed with solubilized thylakoids isolated from *P_35S_:AtCGL160* plants (Fig. 6 A,B), (iv) AtCGL160 co-migrates with a large complex after DSP-mediated crosslinking (Fig. 6C) and (v) AtCGL160N interacts with CF_1_-β in yeast two-hybrid experiments (Fig. 7).

A role for AtCGL160 in the incorporation of CF_1_ into the holocomplex was previously proposed by Fristedt et al. (2015). This assumption was based on the observations that AtCGL160 co-migrated with CF_1_ subcomplexes in BN/SDS-PAGE analyses and could be cross-linked to CF_1_ subunits in wild-type protein samples. However, we detected AtCGL160 predominantly in the monomer fraction in untreated thylakoid preparations in this study (Fig. 6C), as well as in previous work (Rühle et al., 2014) – and co-migration of AtCGL160 with high-molecular-mass complexes was only observed after thylakoid proteins had been crosslinked with DSP (Fig. 6C). Furthermore, the commercially available AtCGL160 antibody (AS12 1853, Agrisera) employed in the study of Fristedt et al. (2015) was found here to cross-react strongly with CF_1_-α or CF_1_-β (Supplemental Fig. 3), which complicates the interpretation of one-dimensional co-migration and crosslinking experiments in the absence of appropriate controls. Therefore, a new antibody was generated that does not cross-react with CF_1_-CF_O_ subunits and thus provides a reliable means of probing the molecular interactions of AtCGL160 (Supplemental Fig. S3).

Besides CGL160, ALB4 – a member of the bacterial ALB3/Oxa1/YidC protein insertase family – was previously proposed to participate in the linkage of a CF_1_ to a CF_O_ assembly module (Benz et al., 2009). Another study provided evidence that ALB4 and its paralog ALB3 physically interact with each other, and show significant functional overlap in the membrane insertion of subunits of the Cyt *b_6_f* complex (Trosch et al., 2015). Moreover, alleles of *ALB4* (*STIC1*) have been identified as suppressors of the chloroplast protein import mutant *tic40* (Bedard et al., 2017), and ALB4/STIC1 and STIC2 were shown to act together in thylakoid protein targeting in a pathway that also involves cpSRP54 and cpFtsY. In our study, we did not identify ALB4/STIC1 in co-IP experiments with anti-AtCGL160 antibodies (Fig. 6, Supplemental Table S1) and amounts of thylakoid-associated CF_1_-β in *atalb4-1* mutants (SALK_136199C) grown under short-day conditions were unaltered (Supplemental Fig. S6). Thus, ALB4/STIC1 does not act in concert with CGL160 in late stages of CF_1_-CF_O_ assembly, but serves as a general thylakoid protein biogenesis factor involved in folding or assembly of a specific subset of transmembrane proteins (Bedard et al., 2017).

### AtCGL160 is critical for chloroplast development in the dark

It has long been thought that the hydrolytic activity of CF_1_-CF_O_ needs to be inactivated in the dark to prevent futile ATP depletion (Ort and Oxborough, 1992). However, analysis of the constitutively redox-activated γ-subunit mutant *gamera*, in which a ‘dark *pmf*’ is maintained, revealed increased stability of photosynthetic complexes upon prolonged darkness, suggesting that a certain degree of ATPase activity may be beneficial during the night (Kohzuma et al., 2017). Concomitantly, several processes have been proposed to depend on the maintenance of a dark *pmf*. These include thylakoid protein transport via the Tat- and Sec-dependent pathways, modulation of protease activity and ion homeostasis in the chloroplast. In this regard, a remarkable influence of AtCGL160 disruption on leaf variegation (Fig. 2) and chloroplast development (Fig. 3) was observed exclusively under short-day conditions. Surprisingly, this phenotype was not detectable in *atcgld11-1* plants with a defect in CF_1_ assembly and reduced amounts of CF_1_-CF_O_ comparable to those in *atcgl160-1* (Fig. 4D). However, the leaf phenotype correlated with the accumulation of a CF_1_ intermediate in the stroma (Fig. 5C). Thus, AtCGL160-mediated CF_1_ recruitment might also be critical in preserving the dark *pmf* at night. Alternatively, stroma-enriched CF_1_ complexes (Fig. 5C) could alter the chloroplast ATP/ADP ratio by excessive hydrolytic activity, and disturb ATP-dependent nocturnal processes that ultimately lead to premature chloroplast degradation (Fig. 3).

### AtCGL160 is a central CF_1_-CF_O_ assembly factor with multiple functions

Assembly of membrane-embedded ATP synthase modules and their subsequent association with F_1_ subcomplexes are critical steps in bacterial and organellar ATP synthase biogenesis, as premature formation of the proton-translocating channel between the c-ring and the a-subunit (equivalent to the ATP9 ring and the ATP6 subunit in mitochondria) can lead to uncontrolled dissipation of the *pmf* (Birkenhäger et al., 1999; Franklin et al., 2004), and only efficient integration of F_1_ triggers ATP production. In this context, molecular aspects of the assembly processes were recently elucidated for bacterial (reviewed in Deckers-Hebestreit, 2013), as well as yeast and human mitochondrial ATP synthases (reviewed in Song et al., 2018). One significant outcome was that, while ATP synthase assembly pathways and the repertoire of auxiliary factors differ among these systems, formation of the proton-translocating unit during the final assembly steps is common to all of them.

Intriguingly, our data revealed a dual involvement of AtCGL160 in CF_1_-CF_O_ assembly, namely in c-ring formation and the recruitment of a CF_1_ intermediate. In fact, these two events were suggested to proceed sequentially in the assembly of bacterial ATP synthases (Deckers-Hebestreit, 2013). Since an *E. coli* strain lacking subunit δ accumulates a c_10_α_3_β_3_γε subcomplex, it is assumed that cytoplasmic F_1_ first binds to the c_10_ ring, and c_10_α_3_β_3_γε associates with the ab_2_ module in a δ-dependent manner in the final assembly step (Hilbers et al., 2013). By analogy with the bacterial assembly pathway, AtCGL160 may facilitate the integration of a stator assembly module into the holo-complex. Indeed, the interaction of AtCGL160C with CF_O_-b has been demonstrated in split-ubiquitin assays (Rühle et al., 2014). Moreover, CF_O_-a was less highly enriched in co-IP analyses than other CF_1_-CF_O_ subunits (Fig. 6A, B), which might argue for the release of AtCGL160 after functional incorporation of CF_O_-a in the final steps of CF_1_-CF_O_ assembly. In this scenario, AtCGL160 could act as a placeholder to prevent the premature formation of proton-translocating intermediates. A similar function has been described for the INA complex in yeast mitochondria, which binds to the c-ring, but also to a distinct assembly intermediate consisting of ATP6, ATP8, ATP10, ATP23, peripheral stalk subunits and the F_1_ domain (Naumenko et al., 2017). This ensures that the c-ring and subunit ATP6 are assembled into the proton-conducting unit in a controlled manner. However, due to a generally low turnover rate of CF_1_-CF_O_ assembly (reviewed in Schöttler et al., 2014) and inefficient detection of distinct thylakoid-integral intermediates, a robust CF_O_ assembly map is still lacking, and ‘true’ stator-containing assembly modules have not been described so far.

Nevertheless, a straightforward assembly mechanism for the recruitment of CF_1_ can be derived from our study. After AtCGL160-assisted ring formation (Rühle et al., 2014), the stromally oriented AtCGL160N (Fig. 1) binds to a CF_1_ intermediate consisting of α_3_β_3_γε but not subunit δ (Fig. 5C, Supplemental Fig. S4). Recruitment is mediated through interaction of AtCGL160_29-105_ with subunit CF_1_-β; thus, the phosphorylatable AtCGL160 segment is dispensable for the interaction (Fig. 7). Since AtCGL160 can be cross-linked to high-molecular-mass complexes that are larger than CF_1_ (Fig. 6C), AtCGL160 might remain attached to a putative c_14_α_3_β_3_γε or bb’c_14_α_3_β_3_γε intermediate. Its release could then be triggered by the incorporation of subunit CF_O_-a or CF_1_-δ in the final assembly steps.

At this stage, we cannot rule out the possibility that AtCGL160N might have regulatory functions beyond CF_1_ recruitment, as it interacts with the thylakoid-proximal domain III of CF_1_-β, which contains the conserved DELSEED motif (Fig. 7B). Several regulatory mechanisms have been elucidated in which the subunit β and the DELSEED motif are implicated. For instance, the autoinhibitory subunit ε interacts with the DELSEED motif in bacteria (Tanigawara et al., 2012; Sobti et al., 2016), whereas in bovine (Cabezon et al., 2003) and yeast mitochondria (Robinson et al., 2013), the small protein IF_1_ inhibits ATPase activity by binding at the α/β interface. In plants, a regulatory mechanism controls CF_1_-CF_O_ activity involving also the DELSEED and an L-shaped, two β-hairpin containing motif with two conserved redox-sensitive cysteines in the CF_1_-γ subunit (Hahn et al., 2018). By analogy with the role of IF_1_, which was shown to inhibit ATPase activity during the assembly of human mitochondrial ATP synthases (He et al., 2018), AtCGL160N may regulate ATPase activity during CF_1_-CF_O_ assembly via an as yet unknown mechanism.

## Methods

### Bioinformatics Sources

Protein and gene sequences were downloaded from the Arabidopsis Information Resource server (TAIR; http://www.arabidopsis.org), Phytozome (https://phytozome.jgi.doe.gov/pz/portal.html) and the National Center for Biotechnology Information server (NCBI; http://www.ncbi.nlm.nih.gov/). Transit peptides were predicted by ChloroP (http://www.cbs.dtu.dk/services/ChloroP/) (Emanuelsson et al., 1999). Structural data was obtained from the PDB homepage (https://www.rcsb.org/) and processed with ChimeraX (https://www.cgl.ucsf.edu/chimerax/) (Pettersen et al., 2021). Multiple sequence alignments were generated with the CLC workbench software (v8.1) and protein features were visualized with Protter (https://wlab.ethz.ch/protter/start/) (Omasits et al., 2014). Co-migration of stromal proteins was examined with the online tool PCom-DB (http://pcomdb.lowtem.hokudai.ac.jp/proteins/top) (Takabayashi et al., 2017). Boxplots were drawn with BoxPlotR (http://shiny.chemgrid.org/boxplotr/) (Spitzer et al., 2014).

### Plant Material and Growth Conditions

T-DNA lines for *atcgl160-1 (*SALK_057229, Col-0 background), *atcgld11-1* (SALK_019326C, Col-0 background) and *atalb4-1* (SALK_136199C) were obtained from the SALK collection (Alonso et al., 2003). Plants were grown on potting soil (A210, Stender, Schermbeck, Germany) under controlled greenhouse conditions (70-90 µmol photons m^-2^ s^-1^, 16/8 h light/dark cycles), or in climate chambers on an 8h light/16h dark cycle for biochemical and physiological analyses. Fertilizer was added to plants grown under greenhouse conditions according to the manufacturer’s recommendations (Osmocote Plus; 15% nitrogen [w/v], 11% [w/v] P_2_O_5_, 13% [w/v] K_2_O, and 2% [w/v] MgO; Scotts, Germany). For domain-specific complementation assays, either the complete coding region of *AtCGL160* (*P_35S_:AtCGL160*) or parts of the CDS corresponding to amino acids 1-206 (*P_35S_:AtCGL160N*) and 207-350 (*P_35S_:AtCGL160C*) were cloned into the binary Gateway vector pB2GW7 (Karimi et al., 2002), placing the genes under control of the 35S CaMV promoter. The transit peptide coding sequence (for amino acids 1-46) was fused to the *AtCGL160C* CDS in the case of the *P_35S_:AtCGL160C* vector. The constructs were first transformed into *Agrobacterium tumefaciens* strain GV3101, and then into *atcgl160-1* plants by the floral-dip method (Clough and Bent, 1998). T1 plants were selected by screening for Basta resistance. Basta positives were screened for equal amounts of the *AtCGL160* transcript by RNA gel-blot hybridization as described below.

### Transmission electron microscopy

Leaf pieces of about 1.5 × 1.0 mm were cut with a new double edge razor blade (Feather, Osaka, Japan) and immediately immersed in fixation buffer (0.1 M sodium phosphate buffer, pH 7.4, 2.5% [v/v] glutaraldehyde, 4% [v/v] formaldehyde) at room temperature. A mild vacuum (about 20 mbar) was applied until the leaf pieces did sink, the fixation buffer replaced with fresh one and the samples fixed overnight at 4 °C. After three 10-min washes in sodium phosphate buffer (pH 7.4), the samples were osmicated with 1% osmium tetroxide and 1.5% potassium ferricyanide in 0.1 M sodium phosphate buffer (pH 7.4) for 60 min at 4°C. The samples were rinsed three times for 10 min each in distilled water and incubated in 1% uranyl acetate (in distilled water) at 4°C overnight. After three washes of 10 min each in distilled water the samples were dehydrated using increasing concentrations of ethanol and infiltrated, with propylene oxide as intermediate solvent, in glycid ether 100 (formerly Epon 812; Serva, Heidelberg, Germany) following standard procedures. Polymerization was carried out for 40 - 48 h at 65 °C. Ultrathin sections (∼60 nm) were cut with a diamond knife (type ultra 35°; Diatome, Biel, Suisse) on an EM UC7 ultramicrotome (Leica Microsystems, Wetzlar, Germany) and mounted on single-slot Pioloform-coated copper grids (Plano, Wetzlar, Germany). The sections were stained using uranylacetate and lead citrate (Reynolds, 1963) and viewed with a JEM-1400 Plus transmission electron microscope (JEOL, Tokyo, Japan) operated at 80kV. Micrographs were taken using a 3.296 × 2.472 pixels charge-coupled device camera (Ruby, JEOL).

### Chl *a* Fluorescence Measurements

*In vivo* Chl a fluorescence of whole plants was measured using an imaging Chl fluorometer (Imaging PAM, Walz, Effeltrich, Germany). Plants were dark-adapted for 20 min and exposed to a pulsed, blue measuring beam (4 Hz, intensity 3, gain 3, damping 2; F_O_) and a saturating light flash (intensity 10) to calculate F_V_/F_M_. If not indicated otherwise, transient NPQ induction was measured at 145 µmol photons m^-2^ s^-1^.

### ECS Measurements

ECS measurements were performed using the Dual-PAM-100 (Walz, Effeltrich, Germany) equipped with a P515/535 emitter-detector module (Schreiber and Klughammer, 2008). The measurement was carried out at 23°C under ambient CO_2_ conditions. Plants grown in short-day conditions for four weeks were light-adapted, and detached leaves were illuminated for at least 10 min with 129 µmol photons m^-2^ s^-1^ red light. After illumination, dark-interval relaxation kinetics (DIRK) were measured in the ms to s range. Values for *pmf* (ECSt), and proton conductivity (gH^+^) were calculated as described (Cruz et al., 2001; Schreiber and Klughammer, 2008). Briefly, the maximum amplitude of the inverse electrochromic band-shift kinetic was measured in the second range, and normalized to a single saturating P515 pulse measured after 4 minutes of dark incubation. For proton conductivity, electrochromic band-shift kinetics were recorded in the millisecond range for 5 consecutive periods of 2 sec, separated by dark intervals of 30 sec. The combined signals were fitted to a single exponential decay function and the reciprocal value of the lifetime was used to estimate the proton conductivity (Kanazawa and Kramer, 2002).

### AtCGL160 Antibody Generation

Rabbit antibodies were generated against AtCGL160 that had been heterologously expressed in *Escherichia coli*, and then purified. To this end, the coding sequence corresponding to AtCGL160_29-206_ was cloned into the pMal-c5x vector (New England Biolabs) and purified on amylose columns (New England Biolabs) according to the manufacturer’s instructions. The protein was injected into rabbits for antibody production (Pineda, Berlin, Germany). To reduce epitope cross-reactions, the antiserum was purified on a column crosslinked with heterologously expressed AtCGL160_29-206_ fused to the glutathione-S-transferase (GST) tag. Purified antibody was employed at a dilution of 1:1000. Signals were detected by enhanced chemiluminescence (Pierce™ ECL Western Blotting Substrate, Thermo Scientific) using an ECL reader system (Fusion FX7; PeqLab, Erlangen, Germany).

### Nucleic Acid Analysis

Total RNA from snap-frozen leaves was extracted with the RNeasy Plant Mini Kit (Qiagen) according to the supplier’s instructions. Samples equivalent to 8 or 20 µg total RNA were fractionated by electrophoresis in formaldehyde-containing agarose gels (1.2%), blotted onto nylon membranes (Hybond-N+, Amersham Bioscience) and fixed by UV radiation (Stratalinker® UV Crosslinker 1800). To control for equal loading, abundant RNAs on nylon membranes were stained with methylene blue solution (0.02% [w/v] methylene blue, 0.3 M sodium acetate pH 5.5). To detect gene-specific transcripts, DNA fragments amplified from cDNA were labelled with radioactive [α-^32^P]dCTP and subsequently used as probes in hybridization experiments (see Supplemental Table S2 for primer information). Signals were detected with the Typhoon Phosphor Imager System (GE Healthcare).

### Protein Analysis

Leaves from 4-week-old plants grown under short-day conditions were harvested shortly after the onset of the light period, and thylakoid membrane-enriched samples were isolated according to Rühle et al. (2014). Crosslinking of thylakoids was performed by incubation with 2.5 mM dithiobis(succinimidyl propionate) (DSP, Thermo Scientific). After incubation for 20 min on ice, crosslinking was quenched with 60 mM Tris/HCl (pH 7.5). Chl concentrations were determined as described in Porra et al. (1989). For immunotitrations, thylakoid membrane pellets were resuspended in loading buffer (100 mM Tris/HCl pH 6.8, 50 mM dithiothreitol [DTT], 8% [w/v] SDS, 24% [w/v] glycerol and 0.02% [w/v] bromophenol blue). Denaturation for 5 min at 70°C and protein fractionation on Tricine-SDS-PAGE gels (10% gels supplemented with 4M Urea) was carried out according to Schägger (2006). Immunodetections were performed as described below. Sample preparation for BN-PAGE was performed with freshly prepared thylakoids as described in Peng et al. (2008). First, membranes were washed twice in wash buffer (20% glycerol, 25 mM BisTris/HCl pH 7.0). Then, samples were treated with wash buffer containing 1% (w/v) n-dodecyl β-D-maltoside and adjusted to 1 ml mg^-1^ Chl for 10 min on ice. After centrifugation (16,000*g*, 20 min, 4°C), supernatants were supplemented with 1/10 volume of BN sample buffer (100 mM BisTris/HCl pH 7.0, 750 mM ε-aminocaproic acid, 5% [w/v] Coomassie G-250). BN-PAGE gels (4-12% gradient) were prepared as described in Schägger et al. (1994). Solubilized samples corresponding to 60 µg Chl were loaded per lane and gels were run at 4°C overnight. To separate complexes into their subunits, BN-PAGE strips were treated with denaturing buffer (0.2 M Na_2_CO_3_, 5% [w/v] SDS, 50 mM DTT) for 30 min at room temperature and loaded on Tricine-SDS-PAGE gels. Gels were subsequently subjected to immunoblot analysis with antibodies against CF_1_-CF_O_ subunits and AtCGL160, as described below.

For analysis of the stromal CF_1_ intermediate, intact chloroplasts from 4-week-old plants were isolated according to Rühle et al. (2021). After lysis in 25 mM HEPES/KOH (pH 7.5) containing 5 mM MgCl_2_ for 30 min on ice, the stromal fraction was separated from membranes by centrifugation at 35,000*g* for 30 min (4 °C). Protein concentration was measured using the Bradford Protein Assay (Bio-Rad). Stromal BN analysis was performed according to Reiter et al. (2020). In brief, chloroplast-enriched pellets were resuspended in BN washing buffer and mechanically disrupted by passage through an 0.45-mm syringe. The stromal fraction was separated from membranes by centrifugation at 35,000*g* for 30 min (at 4°C). 100 µg of total soluble protein was mixed with 1/10 volume of BN sample buffer before fractionation in the first dimension as described above.

### Immunoblot Analyses

Proteins fractionated by gel electrophoresis were transferred to polyvinylidene difluoride membranes (PVDF) (Immobilon®-P, Millipore) using a semi-dry blotting system (Biorad) as described in the supplier’s instructions. After blocking with TBST (10 mM Tris/HCl pH 8.0, 150 mM NaCl and 0.1% [v/v] Tween-20) supplemented with 3% (w/v) skim milk powder, the membranes were incubated with antibodies at 4°C overnight. Antibodies used in this study were obtained from Agrisera (CF_1_-β: AS05 085, 1:5000; CF_1_-γ: AS08 312, 1:5000; CF_O_-b: AS10 1604, 1:5000; CF_O_-c: AS09 591, 1:3000; and AtCGL160: AS12 1853, 1:1000).

### Yeast-two-Hybrid Experiments

Yeast two-hybrid assays were carried out using the Matchmaker Two-Hybrid System Kit (Clontech). The *AtCGL160* CDS without the signal peptide (see Supplemental Table S2 for primer information) was cloned into the bait vector pGBKT7 (BD-AtCGL160), whereas the coding sequences of CF_1_-α, -β, -γ, -δ, -ε, the soluble domains of CF_O_-b (51-184 aa) and b' (109-219 aa), AtCGL160 and the CF_1_ assembly factor AtCGLD11 were cloned into the prey vector pGADT7 (named AD-CF_1_-α, AD-CF_1_-β, AD-CF_1_-γ, AD-CF_1_-δ, AD-CF_1_-ε, AD-CF_O_-b, AD-CF_O_-b', AD-AtCGL160 and AD-AtCGLD11). As in the case of AtCGL160, signal peptide sequences were omitted from the nucleus-encoded subunits CF_1_-γ, CF_1_-δ, CF_O_-b' and AtCGLD11. For binding-domain analysis of CF_1_-β, the respective CDS was sub-divided into three parts, according to Groth and Pohl (2001) and cloned into pGADT7. In the case of AtCGL160N binding-site analysis, sequences coding for 29-74, 75-105, 106-134, 135-160 and 161-206 aa were deleted from the BD-AtCGL160 vector using a site-directed mutagenesis kit (NEB). Primers are listed in Supplemental Table S2. Bait and prey vectors were co-transformed into AH109 yeast strains (Clontech) following manufacturer’s instructions. Co-transformants were selected on synthetic dropout (SD) medium (Clontech) lacking leucine and tryptophan (-LT). In order to identify protein interactions, double transformants were grown on SD medium lacking leucine, tryptophan, histidine, and adenine (-LTHA).

### Co-immunoprecipitation

Freshly extracted thylakoids corresponding to ~10 mg chlorophyll were resuspended in 500 µl extraction buffer (50 mM Tris/HCl pH 7.5, 150 mM NaCl, 1 mM MgCl_2_, 5% [w/v] glycerol, 1% [v/v] Nonidet P40 [NP40], 0.2 mM phenylmethylsulfonyl fluoride [PMSF]) and solubilized for 30 min on ice. After centrifugation at 35,000*g* for 30 min and 4°C, the supernatant was added to 20 µl Dynabeads (Thermo Scientific), equilibrated with equilibration buffer (50 mM Tris/HCl pH 7.5, 150 mM NaCl, 5% [w/v] glycerol, 0.05% [v/v] NP40) and labelled with AtCGL160 antibody according to the manufacturer's instructions. The suspension was incubated with rotation for 3 h at 4°C, washed three times with equilibration buffer, and twice with the same buffer but omitting NP40. Proteins were eluted with 100 µl 0.1 M glycine pH 2.0 for 10 min and neutralized with 100 µl 0.1 M ammonium bicarbonate. After treatment with 10 µl of 45 mM DTT and 10 µl of 0.1 M iodoacetamide, samples were digested with 1.5 µg of trypsin at 37°C overnight. Peptides were desalted with home-made C18 stage tips (Rappsilber et al., 2003), vacuum-dried to near dryness and stored at –80°C. LC MS/MS run and data analysis were performed as described in Reiter et al. (2020).

## Author Contributions

B.R. and T.R. designed research. B.R., L.R., G.M., S.G. and T.R. carried out experiments. B.R., D.L. and T.R. prepared the article. T.R. supervised the whole study.

## Acknowledgments

We thank Tim Scheibenbogen, Michael Berger and Tanja Neufeld for technical assistance with Yeast-two-hybrid experiments, Tim Dreißig for technical assistance with heterologous expression of AtCGL160N, and Paul Hardy for critical comments on the manuscript.

## Supplemental tables

**Supplemental Table S1.** AtCGL160 co-immunoprecipitation experiments. Differential enriched proteins in *P_35S_:AtCGL160* versus *P_35S_:AtCGL160C* samples sorted by log_2_ fold change (-log_10_ *P*-value > 1.5). Nucleus-encoded genes are written in capital letters.

| Protein IDs (Uniprot) | Gene names | log<sub>2</sub> FC | -log<sub>10</sub> p-value |
| --- | --- | --- | --- |
| O82279 | *AtCGL160* | 6.495 | 4.644 |
| P09468 | *atpE* | 6.38 | 2.301 |
| Q42139 | *ATPG* | 5.231 | 4.747 |
| P56760 | *atpH* | 4.886 | 3.176 |
| Q9SSS9 | *ATPD* | 4.672 | 4.523 |
| P19366 | *atpB* | 4.437 | 4.459 |
| P56759 | *atpF* | 4.399 | 5.913 |
| P56757 | *atpA* | 4.235 | 6.41 |
| Q01908 | *ATPC1* | 4.156 | 4.772 |
| Q2HIU0 | *At3g15110* | 3.333 | 3.508 |
| P56758 | *atpI* | 2.799 | 4.104 |
| O49445 | *LECRK72* | 2.346 | 3.115 |
| Q67XC4 | *TBL40* | 2.268 | 2.402 |
| Q8LCQ4 | *LHCA6* | 2.082 | 3.71 |
| A0A1P8B288, Q39099 | *XTH4* | 1.983 | 4.126 |
| Q41963 | *TIP1-2* | 1.907 | 2.821 |
| O22957 | *At2g34040* | 1.632 | 2.829 |
| Q9SRL2, Q9M9X0, F4J8G2, Q9SRL7, Q9S9U3 | *RLP32, RLP33, RLP34, RLP35, RLP53* | 1.564 | 3.162 |
| P38418, A0A1I9LPH1 | *LOX2* | 1.544 | 2.047 |
| Q8LBV4 | *At1g78140* | 1.515 | 2.171 |
| F4IUJ0, F4IUI9 | *At2g26340* | 1.451 | 3.021 |
| Q9SF53, A0A1I9LSB4, Q9M3D2 | *RPL35A, RPL35C* | 1.254 | 2.619 |
| A0A1P8B6D0, Q9SUI4 | *PSAL* | 1.193 | 2.938 |
| Q9FFW9, F4KBJ3 | *At5g38520* | 1.136 | 3.084 |
| Q96242 | *CYP74A* | 1.078 | 2.371 |
| Q9SYW8, F4K8I1 | *Lhca2* | 0.941 | 2.356 |
| Q9SR92 | *STR10* | 0.839 | 2.913 |
| P56777 | *psbB* | 0.807 | 2.072 |
| Q9LHA6 | *At3g28220* | 0.731 | 2.315 |
| Q9S7N7 | *PSAG* | 0.575 | 2.157 |

**Supplemental Table S2.**
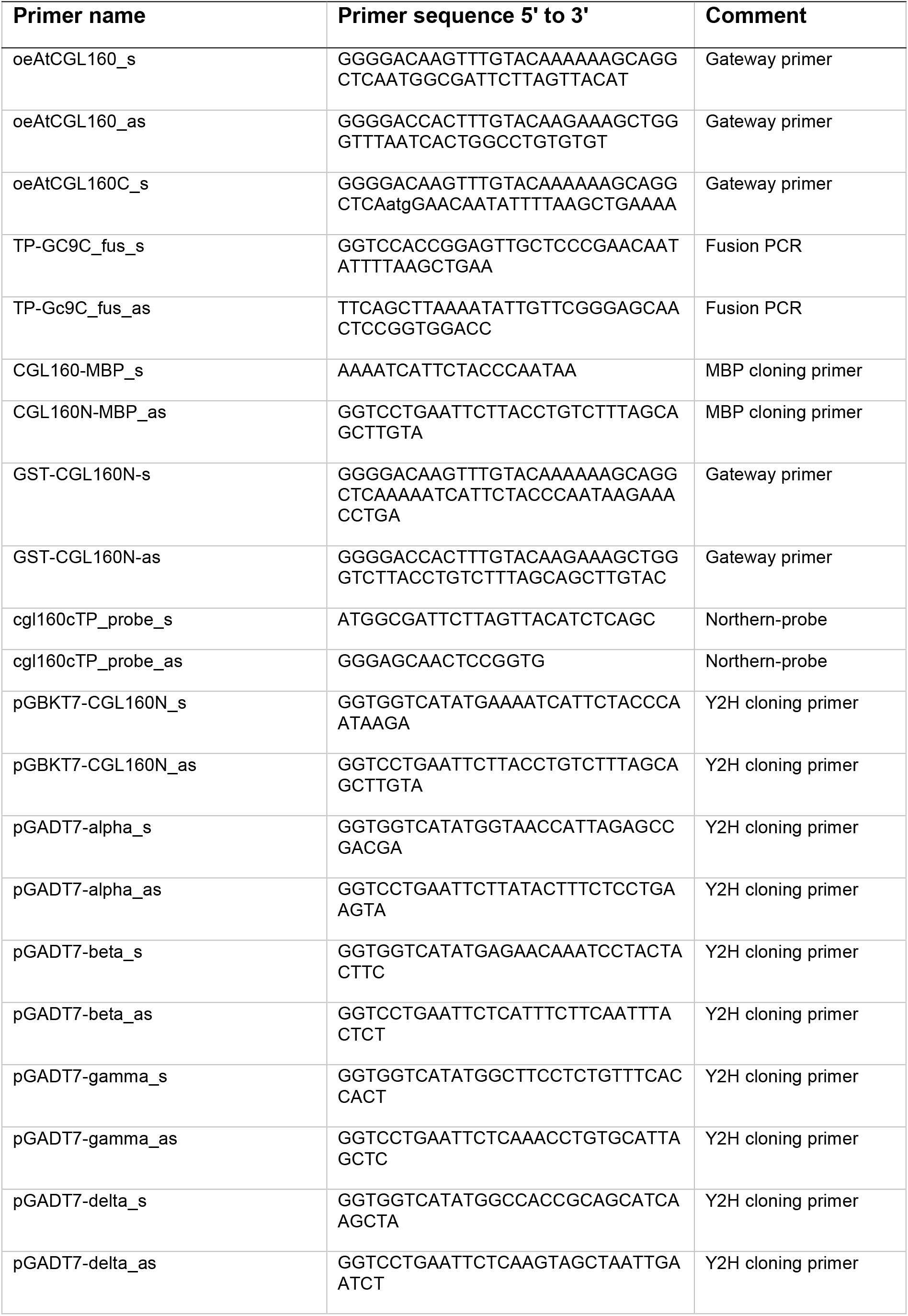
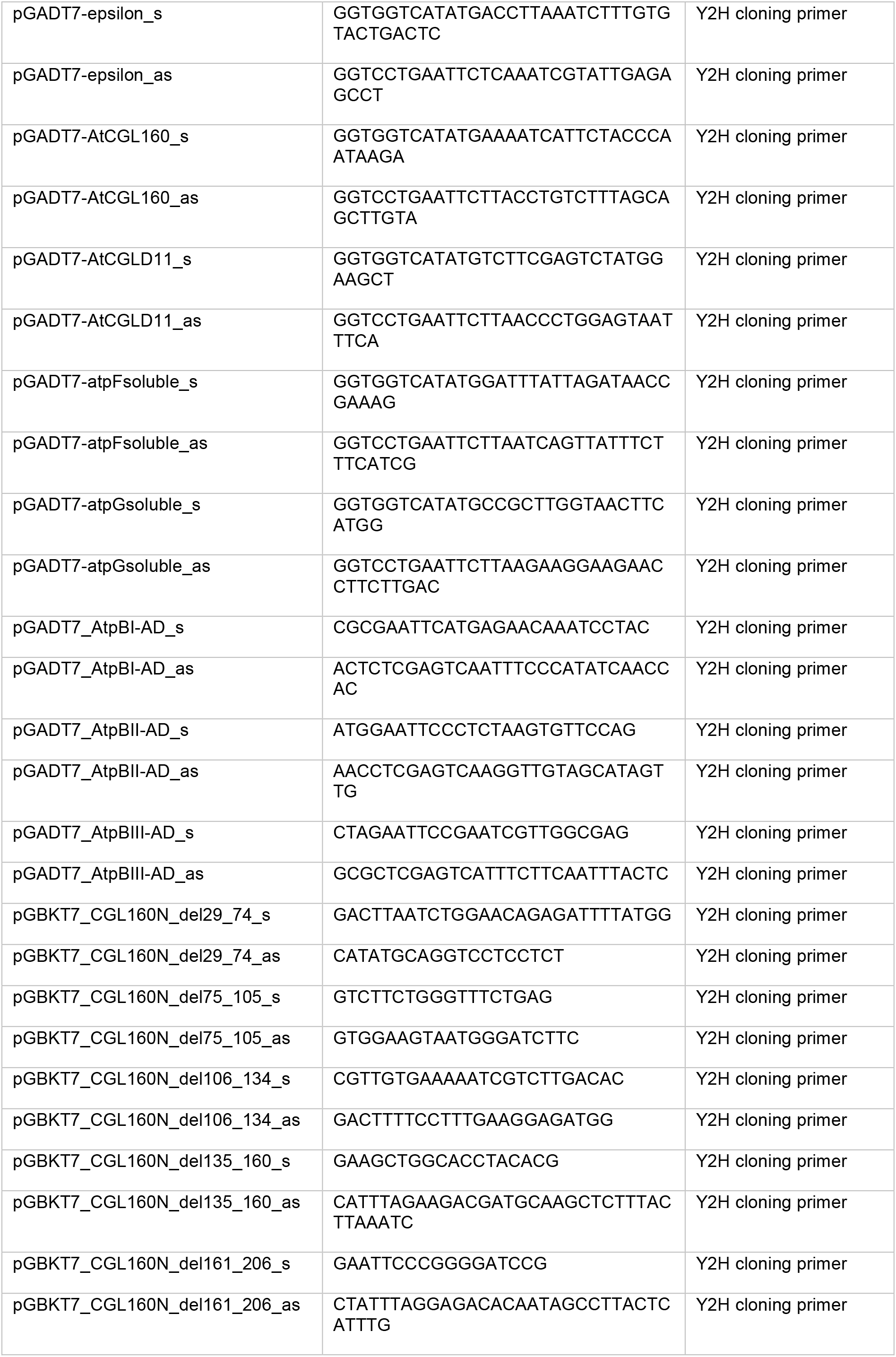
Primers used in this study.

### Supplemental Figures

**Supplemental Figure S1.**
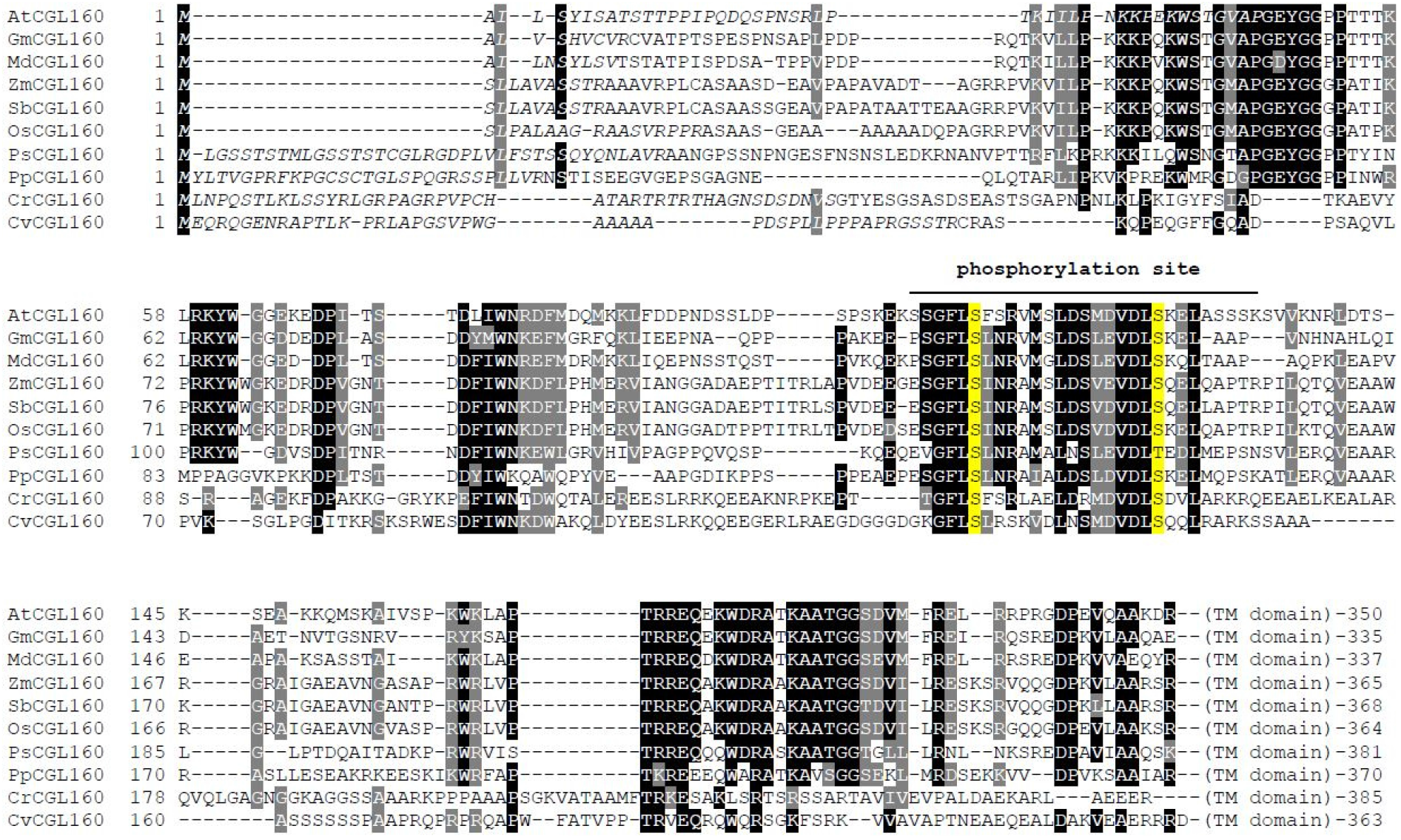
Multiple alignment of the N-terminal portions of CGL160 sequences identified in species belonging to the green lineage. Chloroplast transit peptides predicted by ChloroP are depicted in italics. Similar and identical amino acids conserved in 70% of the sequences are highlighted in grey and black, respectively. The region that includes several identified phosphopeptides in AtCGL160 is indicated and two conserved S/T residues are shown in yellow. Note that CGL160 transmembrane (TM) domains were omitted from the alignment. Sequence identifiers for CGL160 homologs are as follows: *Arabidopsis thaliana* (AtCGL160, NP_565711), *Glycine max* (GmCGL160, XP_006582279.1), *Malus domestica* (MdCGL160, XP_008353735.1), *Zea mays* (ZmCGL160, NP_001170362.2), *Sorghum bicolor* (SmCGL160, XP_021312638.1), *Oryza sativa* Japonica group (OsCGL160, XP_015619276.1), *Picea sitchensis* (PsCGL160, ABR16992.1), *Physcomitrella patens* (PpCGL160, XP_024381807.1), *Chlamydomonas reinhardtii* (CrCGL160, XP_001690237.1) and *Chlorella variabilis* (CvCGL160, XP_005844436.1).

**Supplemental Figure S2.**
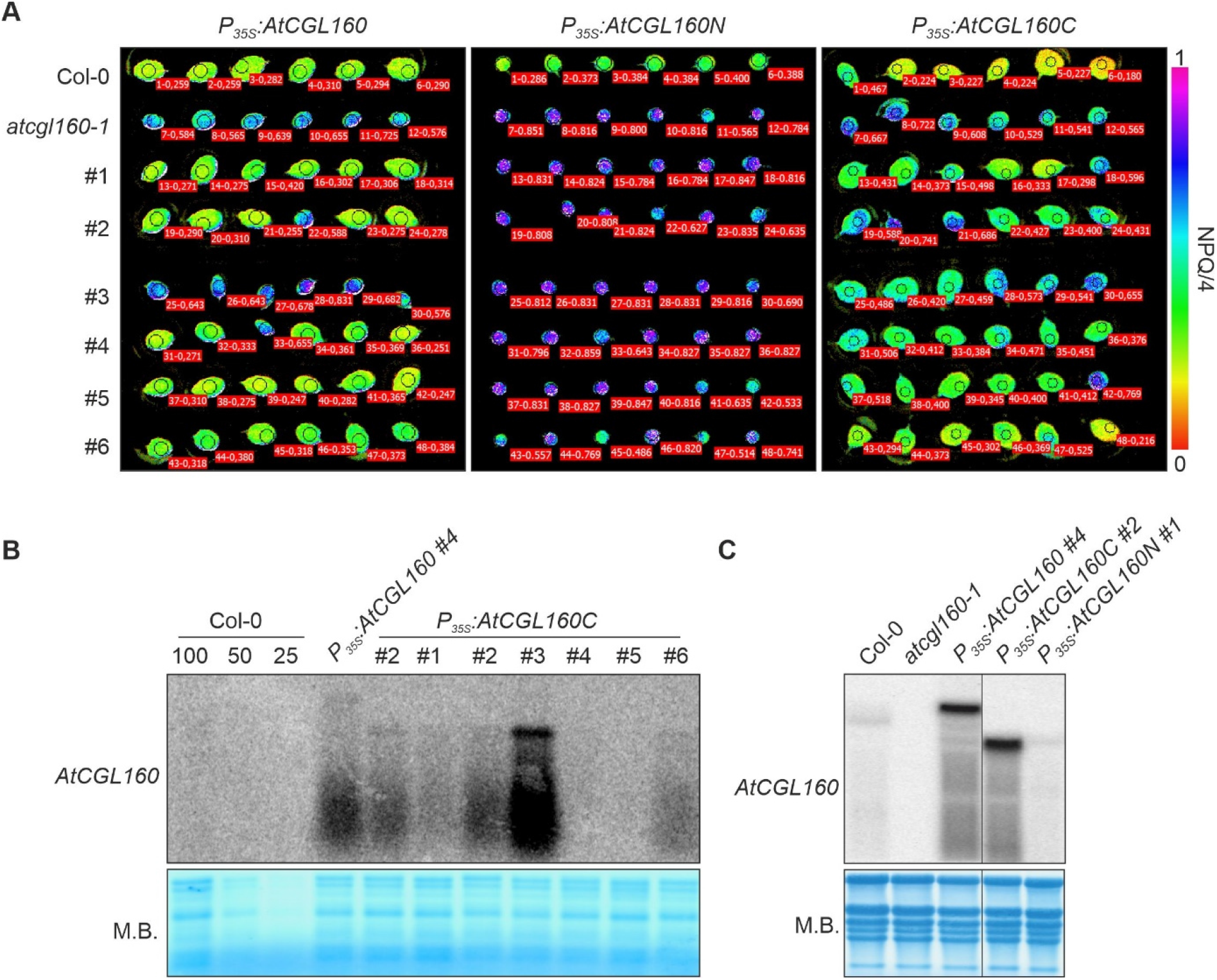
Screening of *P_35S_:AtCGL160, P_35S_:AtCGL160N* and *P_35S_:AtCGL160C* plants. **A**, After transformation of *atcgl160-1*, T2 offspring of independent T1 plants (#1-#6) were examined using an Imaging-PAM (Walz, Effeltrich, Germany) system. Non-photochemical quenching (NPQ/4) was measured in light induction experiments on detached leaves after 8 min of irradiation at 100 µmol photons m^-2^ s^-1^, and is indicated on a false-color scale from 0 to 1. Col-0 and *atcgl160-1* leaves served as controls. *P_35S_:AtCGL160* lines #1, #2, #4, #5 and #6 rescued the *atcgl160-1* phenotype. Transformation of *atcgl160-1* plants with the *P_35S_:AtCGL160N* and *P_35S_:AtCGL160C* constructs resulted in no complementation and partial complementation, respectively. **B**, *AtCGL160* transcript levels in *P_35S_:AtCGL160C* plants determined by Northern analysis. Note that RNA samples (8 µg) of line #2 were loaded twice for direct comparison of transcript levels with line #1 (*P_35S_:AtCGL160*). **C**, Northern analyses of selected, homozygous lines (T3 generation). Total RNA (20 µg) from 4-week-old Col-0, *atcgl160-1*, *P_35S_:AtCGL160, P_35S_:AtCGL160N* and *P_35S_:AtCGL160C* plants was size-fractionated on a denaturing formaldehyde gel and blotted onto a nylon membrane. Hybridization was carried out with a radioactive probe specific for the *AtCGL160* chloroplast transit-peptide coding region. Line #4 (*P_35S_:AtCGL160)* and line #2 (*P_35S_:AtCGL160C)* were selected for further experiments due to their similar transcript levels. Methylene blue (M.B.) staining of the nylon membrane served as an RNA loading control in **B** and **C**.

**Supplemental Figure S3.**
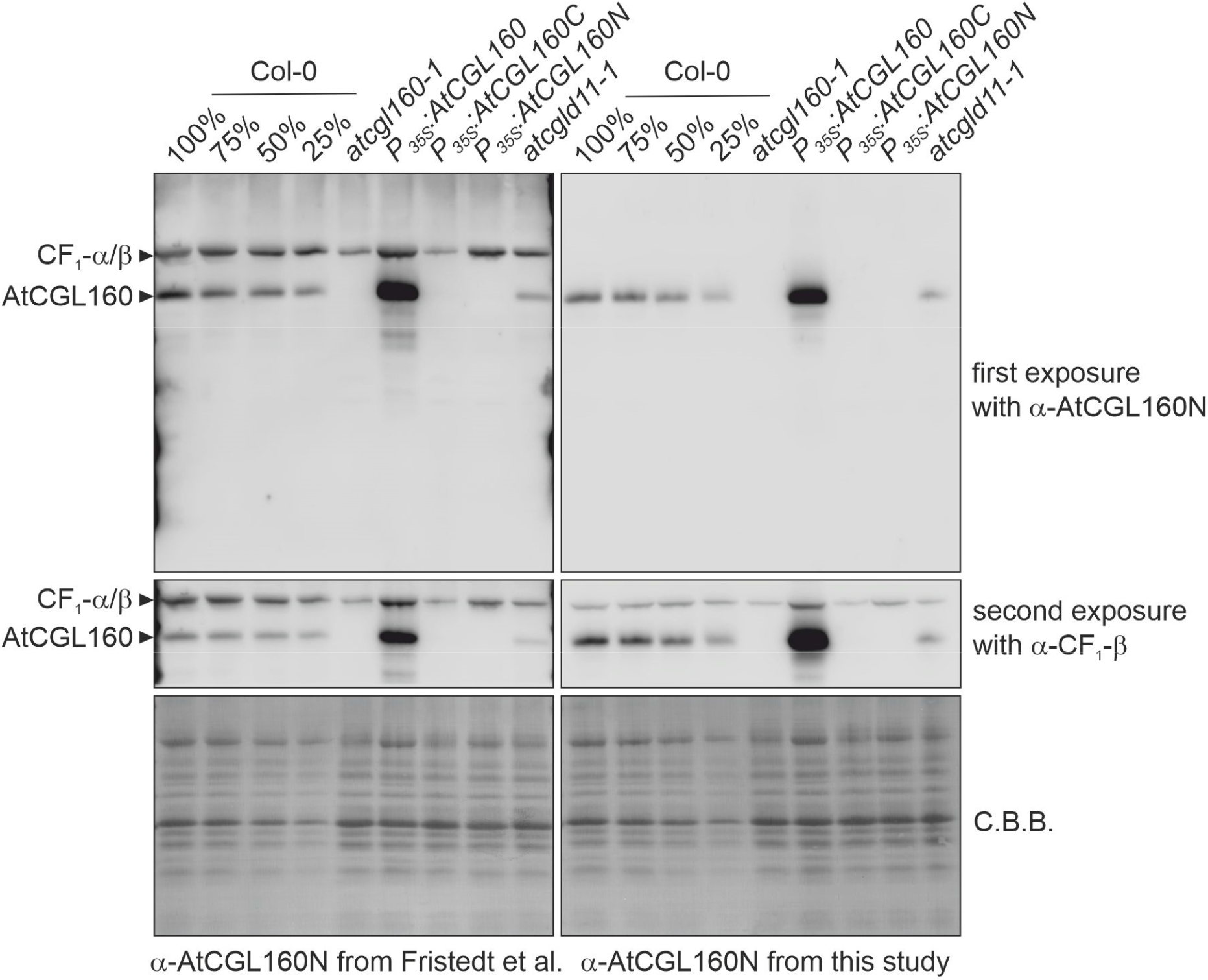
Immunodetection of AtCGL160 in Col-0, *atcgl160-1*, *P_35S_:AtCGL160, P_35S_:AtCGL160N, P_35S_:AtCGL160C* and *atcgld11-1* plants. Thylakoid proteins were separated by denaturing SDS-PAGE and blotted onto PVDF membranes. Membranes were first probed with antibodies against AtCGL160N. After signal detection, membranes were re-probed with an antibody against CF_1_-β. Coomassie brilliant blue staining (C.B.B.) of PVDF membranes is shown as a loading control. On the left, immunodetection analyses are shown for an AtCGL160 antibody (AS12 1853) which is commercially available from Agrisera and was employed in Fristedt et al. (2015). A side-by-side comparison with the newly generated antibody against the N-terminal part of AtCGL160 is provided in the right panel. Note that antibody AS12 1853 from Agrisera binds nonspecifically to CF_1_-α or CF_1_-β and was therefore not considered for use in co-immunoprecipitation, cross-linking or 2D native/SDS-PAGE experiments.

**Supplemental Figure S4.**
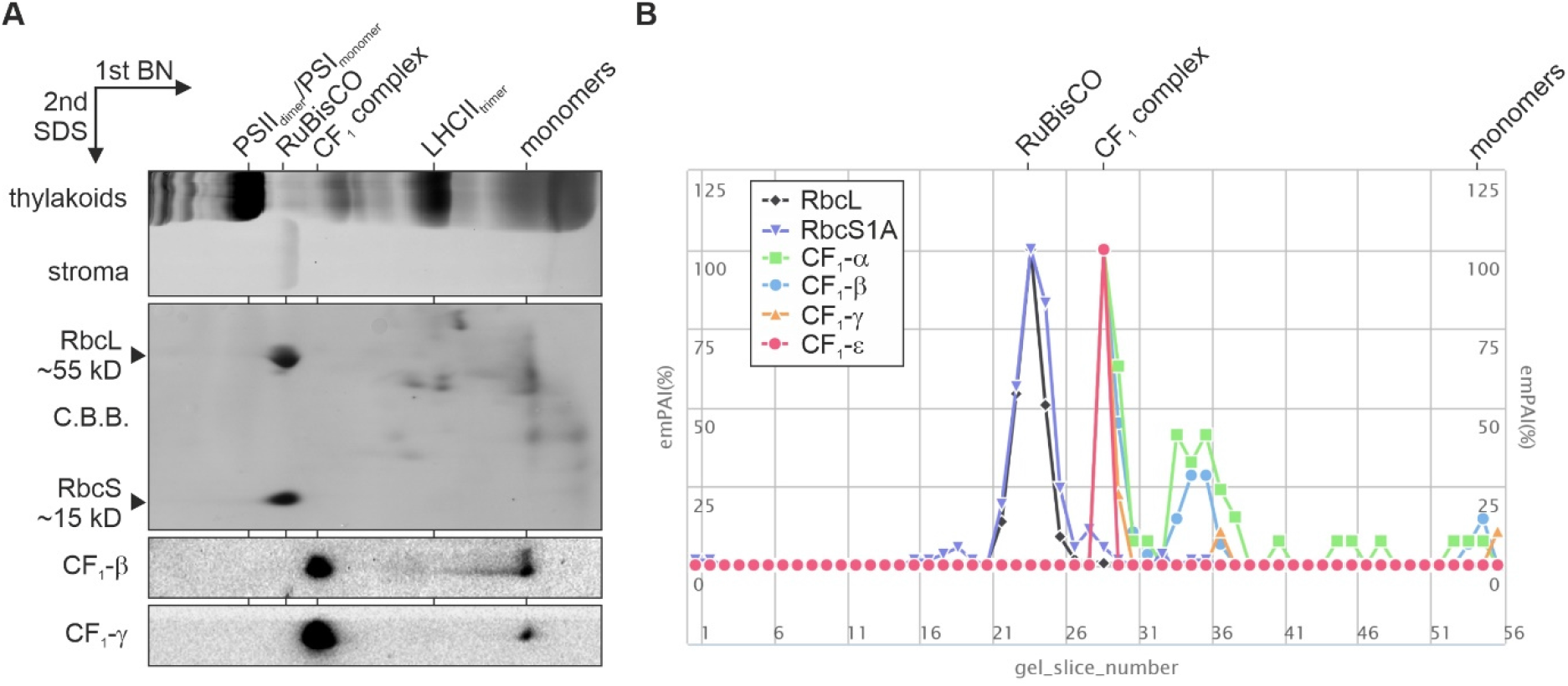
Characterization of the stromal CF_1_ complex in *atcgl160-1* plants. **A**, A stromal protein extract of *atcgl160-1* plants was subjected to 2D gel electrophoresis (Blue Native- and SDS-PAGE) and immunodetection of CF_1_-β and CF_1_-γ. Coomassie brilliant blue (G-250) staining of the PVDF membrane after transfer visualized abundant stromal complexes such as RuBisCO, which is composed of RbcL and RbcS. Prominent thylakoid complexes of *P_35S_:AtCGL160C* plants served as molecular mass standards. **B**, Composition of the stromal CF_1_ sub-complex in Arabidopsis according to the Protein Co-migration Database for photosynthetic organisms (PCom-DB, http://pcomdb.lowtem.hokudai.ac.jp/proteins/top). Co-migration of RbcL (black diamonds) and RbcS (purple triangles) is provided for better comparison between PCom-DB results and the 2D gel analyses presented in panel **A**. Subunit content is quantified according to the exponentially modified protein abundance index (emPAI) method and is normalized for each individual subunit to the maximal emPAI identified in a gel slice (Ishihama et al., 2005). The maximal RuBisCO and CF_1_ content was detected in gel slices 24 and 29, respectively. Note that CF_1_-δ was not identified in the stromal CF_1_ subcomplex.

**Supplemental Figure S5.**
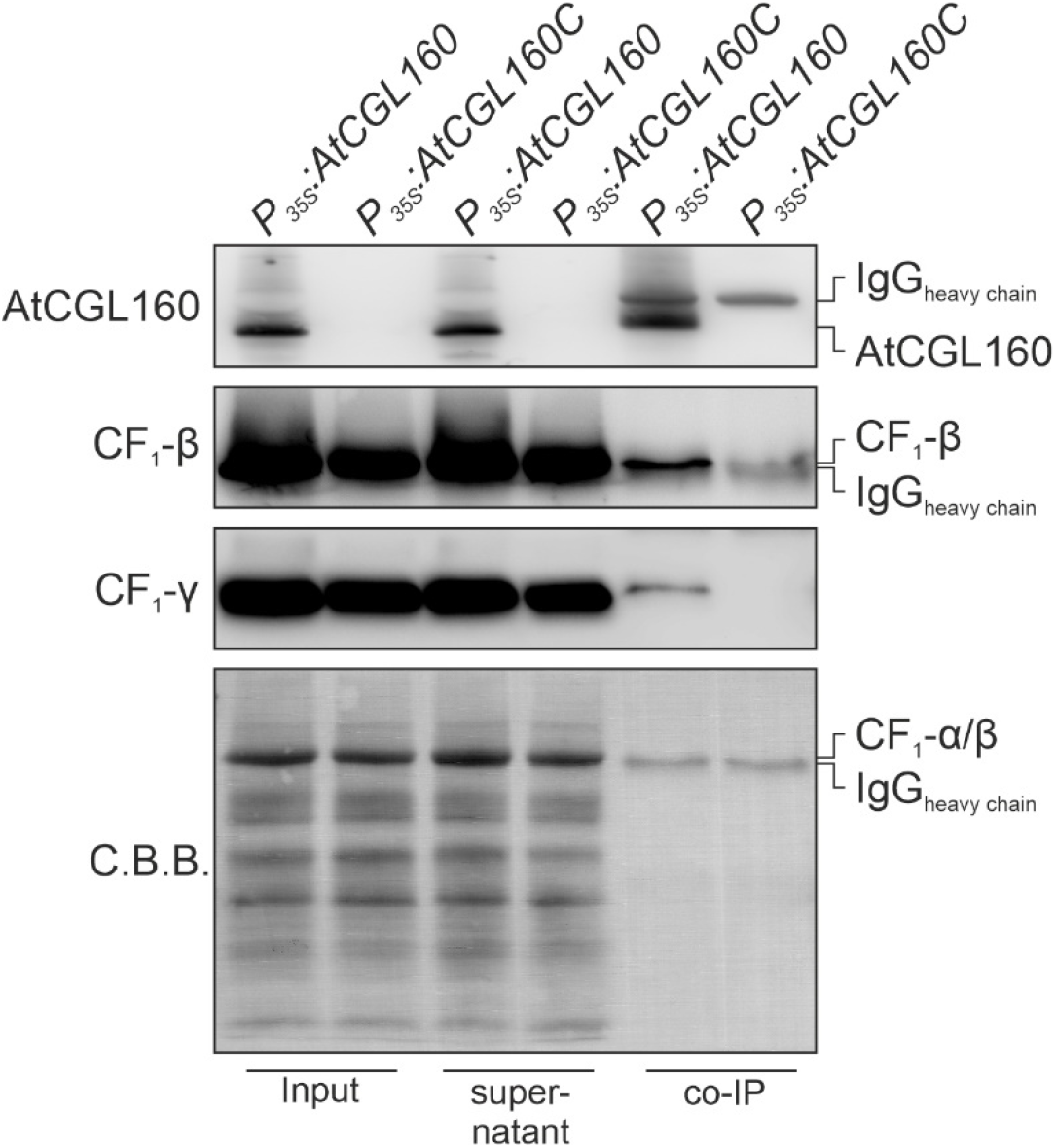
Immunoblot analysis of AtCGL160 co-immunoprecipitation assays. A, Co-immunoprecipitation with NP40-solubilized thylakoids of oeAtCGL160 and oeAtCGL160C was repeated using reduced amounts of the AtCGL160 antibody. Protein A-coupled magnetic beads (Dynabeads, Thermo) coated with AtCGL160 antibody and co-immunoprecipated proteins (IP) were boiled in SDS loading buffer, separated by denaturing SDS-PAGE and blotted onto PVDF membranes. Samples of NP40-solubilized thylakoids before (Input) and after (Flow) incubation with AtCGL160 antibody were loaded as controls. Membranes were probed separately with antibodies against AtCGL160N and CF_1_-β/CF_1_-γ. The positions of the heavy chain of the AtCGL160 antibody are indicated (IgG). Coomassie brilliant blue staining (C.B.B.) is shown as loading control, and the positions of CF_1_-α/β are indicated.

**Supplemental Figure S6.**
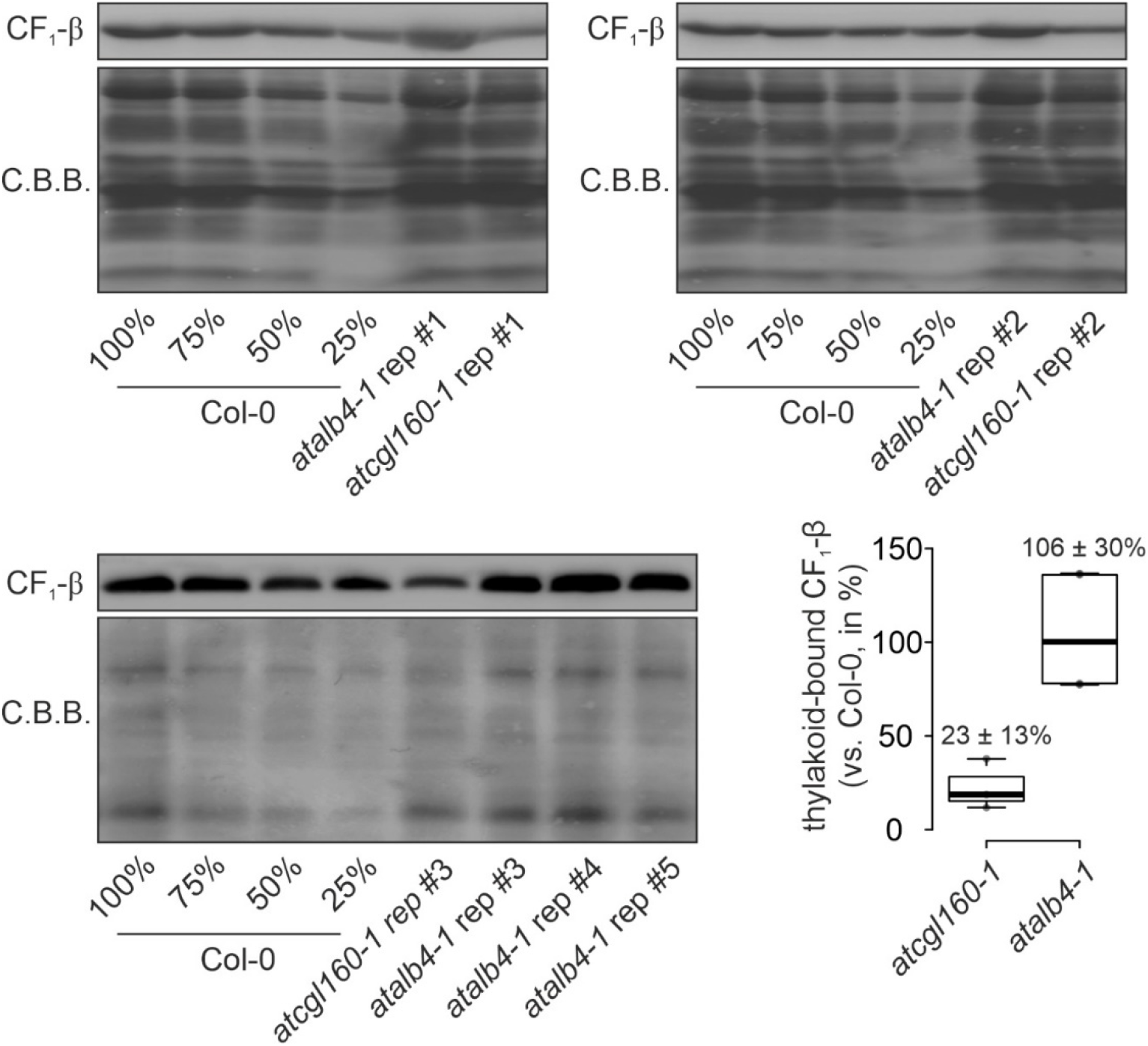
Quantification of thylakoid-bound CF_1_-β subunits in *atalb4-1* Arabidopsis mutant lines. Thylakoid proteins were isolated from Col-0, *atcgl160-1* and *atalb4-1* (SALK_136199C) plants grown under short-day conditions, fractionated on SDS-PAGE, transferred to PVDF membranes and probed with CF_1_-β antibodies. Membranes were stained with Coomassie brilliant blue G-250 (C.B.B.) as loading control. Signals were quantified relative to signals detected in the wild-type sample using the Bio-1D software (version 15.03, Vilber Lourmat, Eberhardzell, Germany) and are provided as percentages. Horizontal lines represent the median, and boxes indicate the 25th and 75th percentiles. Whiskers extend the interquartile range by a factor of 1.5×. Means ± standard deviations are provided above the boxes. Quantification is based on three and five replicates (rep) for *atcgl160-1* and *atalb4-1* samples, respectively.

